# Periventricular Diffusivity Reflects APOE4-modulated Amyloid Accumulation and Cognitive Impairment in Alzheimer’s Continuum

**DOI:** 10.1101/2025.04.28.651021

**Authors:** Chang-Le Chen, Sang Joon Son, Noah Schweitzer, Hecheng Jin, Jinghang Li, Linghai Wang, Shaolin Yang, Chang Hyung Hong, Hyun Woong Roh, Bumhee Park, Jin Wook Choi, Young-Sil An, Sang Woon Seo, Yong Hyuk Cho, Sunhwa Hong, You Jin Nam, Davneet S. Minhas, Charles M. Laymon, George D. Stetten, Dana L. Tudorascu, Howard J. Aizenstein, Minjie Wu, Mayo Clinic Study of Aging

**Author notes:** These authors contributed equally to this work (Chang-Le Chen, Sang Joon Son). Correspondences to: Minjie Wu, PhD., Department of Psychiatry, University of Pittsburgh School of Medicine, 3501 Forbes Ave, Pittsburgh, PA 15213, USA.

## Abstract

**Background:** Altered glymphatic-related fluid dynamics are increasingly recognized as a key feature of Alzheimer’s disease (AD). We generalized an established diffusion imaging technique to estimate periventricular diffusivity (PVeD), hypothesizing that fast diffusion signals in the periventricular region can reflect amyloid-beta (Aβ) deposition across the Alzheimer’s continuum.

**Methods:** Participants from two multi-site cohorts (n = 440 and 414), comprising cognitively unimpaired individuals, those with mild cognitive impairment, and patients with AD, were included. We tested and validated the association of PVeD with Aβ burden and core AD characteristics.

**Results:** Lower PVeD was extensively associated with greater Aβ burden, neurodegeneration, cognitive impairment, and clinical severity. Importantly, the relationship between PVeD and Aβ burden was significantly modulated by APOE4 status, with APOE4 carriers showing a stronger negative association. Baseline PVeD also predicted longitudinal cognitive decline.

**Discussion:** These findings suggest that periventricular fast diffusion signals can reflect APOE4-modulated Aβ burden and cognitive decline in AD.

**Research-in-Context:** *Systematic review:* A comprehensive PubMed literature search indicates that fluid movement related to glymphatic activity assessed by diffusion tensor image analysis along the perivascular space (DTI-ALPS) is associated with amyloid-beta deposition in Alzheimer’s disease (AD). However, recent evidence underscores certain limitations of DTI-ALPS, suggesting that it may not fully capture the diffusion processes involved in amyloid clearance. Moreover, no previous studies have investigated the role of APOE4 in modulating the relationship between glymphatic-related fast diffusion signals and amyloid-beta deposition.

*Interpretation:* The transverse diffusion process along the perivenous space in the periventricular region appears to reflect glymphatic-related dysfunction manifested by amyloid-beta deposition. Reduced periventricular diffusivity is associated with greater amyloid burden across the AD continuum. This association is notably enhanced in APOE4 carriers, who exhibit higher amyloid accumulation for a given reduction in the periventricular diffusivity. Besides, periventricular diffusivity is related to other pathological markers of AD, including clinical symptom severity and neurodegeneration, and may also predict subsequent cognitive decline.

*Future directions:* Although diffusion-based neuroimaging metrics hold promise as surrogate imaging biomarkers for glymphatic-related activity, they do not comprehensively capture the complex fluid dynamics such as convective bulk flow within the glymphatic system. By leveraging multimodal neuroimaging techniques and advanced analytic approaches, future research can refine these metrics into more sensitive, non-invasive tools capable of evaluating fluid dynamics related to glymphatic dysfunction in AD.

## 1 Background

Dementia is characterized by progressive impairments in multiple cognitive domains driven by neurodegenerative processes.^1, 2^ A critical focus of investigation is the glymphatic system, a brain-wide network of perivascular pathways facilitating the exchange of cerebrospinal fluid (CSF) and interstitial fluid (ISF) within the brain.^3, 4^ Glymphatic dysfunction can lead to the accumulation of toxic proteins such as amyloid-beta (Aβ) and tau proteins, which contributes to the pathogenesis of neurodegenerative diseases such as Alzheimer’s disease (AD) and other dementias.^2, 3, 5^ The apolipoprotein E ε4 (APOE4) allele, a known major genetic risk factor for AD, has been linked to glymphatic dysfunction, impairing perivascular clearance and promoting Aβ deposition.^6^ The interplay between impaired glymphatic function and Aβ accumulation may create a feedback loop where waste build-up further compromises glymphatic function, leading to greater neuronal damage and cognitive decline.^5^ Thus, elucidating the mechanisms underlying glymphatic dysfunction, particularly glymphatic fluid dynamics (e.g., convection and diffusion), is critical for identifying therapeutic targets that address the causes related to AD.^2, 7^

The original model of the glymphatic system proposed a convective bulk flow of CSF through perivascular spaces (PVS) driven by arterial pulsations, facilitating the exchange with ISF and the clearance of solutes including Aβ.^2, 3, 8–11^ However, accumulating evidence suggests that fluid movement in the brain parenchyma (i.e. interstitial space, ISS) occurs via the combination of convection and diffusion since bulk flow is potentially limited to the ISS.^2, 4, 8, 12–14^ Also, the impairment of CSF-ISF exchange has been linked to the accumulation of Aβ.^6, 14–17^ Current imaging techniques including contrast-enhanced magnetic resonance imaging (MRI) and diffusion tensor imaging (DTI) have been employed to investigate fluid dynamics relevant to CSF-ISF exchange in the brain.^4, 18–22^ While contrast-enhanced MRI visualizes fluid movement using tracers, DTI assesses the diffusion of water molecules associated with ISF dynamics in perivascular spaces.^18, 23, 24^

The diffusion tensor image analysis along the perivascular space (DTI-ALPS) index, calculated as the ratio of diffusivity along PVS to that perpendicular to main directions of white matter tracts within a certain region of interest (ROI) in the periventricular area, has been hypothesized and investigated as a surrogate marker of glymphatic-related integrity.^25–27^ However, DTI-ALPS has received criticism as a glymphatic-specific marker as it does not distinguish Brownian motion of water along perivascular spaces from other sources of water motion such as neighboring white matter fiber tracts and their structure.^28, 29^ Even so, the DTI-ALPS index has shown significant correlations with cognitive decline, Aβ deposition, and neurodegeneration in AD, suggesting DTI-ALPS as a potential marker for studying the disease.^26, 30, 31^ Nevertheless, the accuracy of DTI-ALPS is highly dependent on predefined small ROIs, making it susceptible to inter-rater variation and image registration errors in automatic analysis, especially with severe brain atrophy, thereby insufficiently capturing effective perivascular diffusion signals. Moreover, while a significant negative association between DTI-ALPS and Aβ burden has been reported, the potential influence of the genetic risk factor, APOE4 allele, on this relationship remains unclear.

Building upon the DTI-ALPS framework, to better capture diffusion signals in periventricular regions relevant to ISF dynamics, we hypothesize that the transverse water diffusion signal at the scale of fast diffusion process within periventricular regions may serve as an imaging correlate of the ISF efflux along the perivenous conduits^2^, which is associated with Aβ clearance and potentially links to pathophysiological characteristics of AD (Figure 1). We proposed a method to estimate an apparent transverse portion of periventricular diffusivity (PVeD) as a diffusion proxy. Our method involves quantifying the voxel-wise transverse diffusivity components in regions adjacent to the lateral ventricles covering main deep medullary veins. Furthermore, we hypothesize the APOE4 allele could modulate the association between ISF and Aβ deposition. The objectives of this study are threefold: (1) to develop and validate the effectiveness of PVeD metrics using clinical DTI data; (2) to associate the PVeD metrics with Aβ burden and other clinical characteristics reflecting neuropathological changes in the AD continuum; and (3) to examine the genetic modulation of APOE4 on the relationship between PVeD and Aβ deposition.

**Figure 1.**
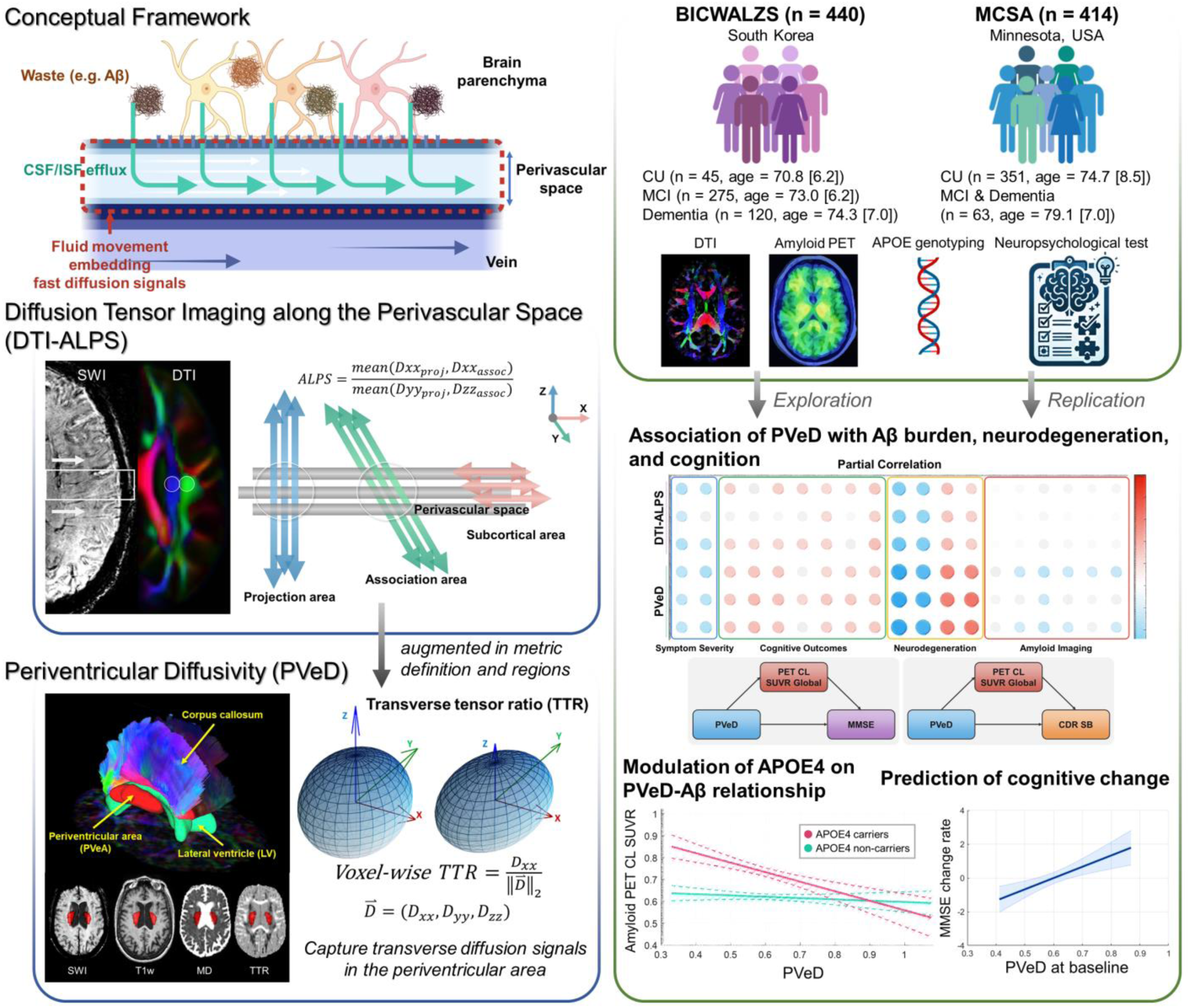
Overview of the study. A conceptual framework of fluid dynamics describes how, within the glymphatic system, solutes such as metabolic waste are propelled toward the perivenous conduit, accompanied by cerebrospinal fluid (CSF) and interstitial fluid (ISF) exchange from the brain parenchyma. This study builds on the method proposed by Taoka et al., who introduced diffusion tensor image (DTI) analysis along the perivascular space (DTI-ALPS) to estimate diffusion-based proxies related to glymphatic integrity given this framework. Rather than targeting predefined regions of interest, we developed an automated approach to delineate the periventricular region using DTI data. We then calculated the transverse tensor ratio (TTR) on a voxel basis to capture the transverse component of fast diffusion signals within the periventricular region, which primarily contains deep medullary veins oriented along the left-right axis. We hypothesize that such periventricular diffusivity (PVeD) can reflect glymphatic-related activities such as amyloid-beta (Aβ) deposition. To test this, we analyzed two multi-site, multi-modal datasets that included cognitively unimpaired (CU) individuals, individuals with mild cognitive impairment (MCI), and patients with AD and other types of dementia for exploration and replication. Our investigation focused on (1) the association of PVeD with Aβ burden, symptom severity, neurodegeneration, and cognitive function, and (2) the modulatory effect of apolipoprotein E ε4 (APOE4) on the association between PVeD and Aβ burden. Additionally, we tested whether the baseline PVeD metrics can predict longitudinal cognitive change. Abbreviations: CDR-SB, Clinical Dementia Rating Sum of Boxes; CL, Centiloid; MD, mean diffusivity; MMSE, Mini-Mental State Examination; PET, positron emission tomography; SWI, susceptibility-weighted imaging; T1w, T1-weighted imaging.

## 2 Methods

### 2.1 Study Cohorts

This study utilized two large-scale, multimodal neuroimaging datasets to test our hypotheses and independently replicate the findings. Specifically, we analyzed data from the BICWALZS and MCSA cohorts, which collectively cover the spectrum of dementia ranging from cognitively unimpaired (CU) individuals to those with clinically diagnosed dementia (Figure 1). These datasets, derived from two distinct national populations, could provide a robust framework for evaluating the link between periventricular diffusion signals and Aβ deposition reflecting the glymphatic-related integrity and clinical characteristics of dementia.

#### 2.1.1 BICWALZS dataset for main analyses

This study leveraged clinical data from the Biobank Innovations for Chronic Cerebrovascular Disease With Alzheimer’s Disease Study (BICWALZS) and the Centre for Convergence Research of Neurological Disorders.^32, 33^ Initiated in 2016 by the Korea Disease Control and Prevention Agency as part of the Korea Biobank Project, BICWALZS supports systematic collection and utilization of human biological specimens and real-world clinical data.^32, 33^ Participants were recruited from memory clinics and a geriatric mental health center with centralized protocols ensuring standardized data collection and harmonization. Clinical diagnoses were established using internationally recognized criteria to identify subjective cognitive decline (SCD), mild cognitive impairment (MCI), Alzheimer’s disease (AD), subcortical vascular dementia (SVaD), etc. Exclusion criteria included major neurological or systemic conditions that could confound findings. The detailed recruitment criteria can be found in Supplementary Material S1. Comprehensive assessments involved symptom severity, neuropsychological testing, amyloid positron emission tomography (PET), APOE genotyping, and brain MRI examination including T1-weighted imaging and diffusion MRI. A longitudinal protocol included annual brief assessments such as Mini-Mental State Examination (MMSE) for the participants and biannual comprehensive evaluations for those with Aβ burden, vascular pathology, APOE4 positivity, or cognitive decline. The BICWALZS study was approved by the Institutional Review Board of Ajou University Hospital (AJOUIRB-SUR-2021-038) and registered in the Korean National Clinical Trial Registry (KCT0003391). Written informed consent was obtained from all participants and caregivers.

#### 2.1.2 MCSA dataset for replication analyses

To validate the findings found in the BICWALZS dataset, we also collected the data from the Mayo Clinic Study of Aging (MCSA), which is a longitudinal population-based cohort study designed to investigate the prevalence, incidence, and risk factors of MCI and dementia among residents of Olmsted County, Minnesota.^34, 35^ The MCSA aims to assess the conversion rates from MCI to dementia or AD, identify risk factors for cognitive decline, and develop predictive models to aid in early detection and prevention strategies. The study cohort consists of individuals mainly aged 50 and older, identified through the Rochester Epidemiology Project. Participants were randomly selected using an age- and sex-stratified sampling frame and underwent a structured baseline evaluation between 2004 and 2020. The assessment included a physician-conducted neurological examination, administration of the Short Test of Mental Status, neuropsychological testing, and structured interviews conducted by researchers to collect demographic data, medical history, and clinical dementia ratings. Standardized diagnostic criteria were applied to determine cognitive status, and vascular risk factors were systematically extracted from electronic health records. In addition to clinical evaluations, a subset of participants underwent multimodal neuroimaging including brain MRI examination and amyloid PET of which imaging data were processed and de-identified for research use.^36^ The neuroimaging data is publicly accessible upon request through the Image & Data Archive run by the Laboratory of Neuro Imaging (https://ida.loni.usc.edu/). All study protocols were approved by the Mayo Clinic Institutional Review Board, and informed consent was obtained from participants or their surrogates.^34, 35^

### 2.2 Neuroimaging Acquisition and Processing

We used DTI data from the BICWALZS and MCSA projects to quantify diffusion signals related to ISF. Other neuroimaging modalities were used to evaluate both neurodegeneration via hippocampal volume (structural MRI) and Aβ deposition (amyloid PET).

#### 2.2.1 MRI imaging

Participants enrolled in the BICWALZS project completed the baseline brain MRI scans on 3T MRI scanners including T1-weighted imaging (in-plane voxel size 0.5-1.0 mm^2^ with slice thickness 1.0-1.3 mm) and single-shell DTI (in-plane voxel size 0.88-1.8 mm^2^ with slice thickness 2.0-3.0 mm, b-value 600-1,000 s/mm^2^). The details of imaging parameters are provided in Supplementary Table 1. T1-weighted images went through image quality assurance procedure, image preprocessing, segmentation, and regional sampling to estimate bilateral hippocampal volume by using CAT12 toolbox (https://neuro-jena.github.io/cat/).^37^ We used multiple anatomical atlases including the AAL3, IBSR, LPBA40, MORI, and Neuromorphometrics built in the CAT12 to obtain unbiased average estimates of hippocampal volume to represent a quantitative measure of neurodegeneration. The hippocampal volume was further adjusted by the total intracranial volume.

Similarly, participants from the MCSA dataset completed the baseline brain MRI scans on 3T MRI scanners including structural MRI (in-plane voxel size 1.0 mm^2^ with slice thickness 1.2 mm) and DTI (in-plane voxel size 1.37 mm^2^ with slice thickness 2.7 mm, b-value 1,000 s/mm^2^). Instead of estimating hippocampal volume using our analytical pipeline, we directly used the estimated hippocampal volume (variable name *SPM12_HPVOL*) and total intracranial volume (variable name *SPM12_TIV*) provided by the MCSA online data repository for analyses.

#### 2.2.2 PET imaging

For the BICWALZS cohort, all participants in this study underwent ^18^F-flutemetamol PET imaging using PET/computed tomography scanners for amyloid imaging. A standardized imaging protocol was implemented across all participants (detailed imaging parameters are provided in Supplementary Table 1). ^18^F-flutemetamol was administered as a single bolus injection (mean dose: 185 MBq) into the antecubital vein. PET acquisition was initiated 90 minutes’ post-injection and continued for 20 minutes, comprising four 5-minute frames. Individual PET scans were co-registered to corresponding MRI scans and subsequently normalized to a standardized T1-weighted MRI template using spatial transformation parameters. ^18^F-flutemetamol retention was quantified using the standard uptake value ratio (SUVR) by which the pons served as the reference region. Global cortical ^18^F-flutemetamol retention was calculated using the automated anatomical labeling (AAL) atlas,^38^ deriving volume-weighted average SUVRs from multiple bilateral cortical ROIs encompassing the frontal, cingulate, temporal, parietal, and occipital lobes. The Centiloid (CL) method was used to convert SUVR values into a standardized, comparable scale for subsequent analyses.^39, 40^

For the MCSA cohort, amyloid PET imaging was performed using ^11^C-Pittsburgh compound B (PiB) according to previously published protocols.^41^ PET images were processed by the MCSA, and the quantification of Aβ burden was performed by calculating the SUVR for each participant using ^11^C-PiB retention. Similarly, we used the global CL SUVR values (variable name *SPM12_PIB_CENTILOID*) provided by the MCSA online data repository for analyses.

### 2.3 DTI Processing

The diffusion weighted images (DWIs) used in this study (both BICWALZS and MCSA datasets) were processed using the DSI studio (version: Chen, March 18 2023, https://dsi-studio.labsolver.org/). The DWIs were first corrected for phase distortion through FSL’s TOPUP if the images acquired with the reverse phase encoding direction were available. This correction was followed by eddy current and head movement corrections using FSL’s EDDY. These FSL correction functions (https://fsl.fmrib.ox.ac.uk/fsl/) were implemented in the integrated preprocessing in the DSI studio. The corrected DWIs were further reconstructed by using *q*-space diffeomorphic reconstruction (QSDR) method,^42^ which is an extension of generalized *q*-sampling imaging approach^43^ that can non-linearly transform the images to the ICBM152 adult template and reconstruct the DWIs in the MNI space. To calculate the diffusivity of interstitial fluid, the DWIs with a b-value of no more than 1,100 s/mm^2^ were used to form the diffusion tensor model. The accuracy of b-table orientation was verified by comparing the fiber orientations with those in a population-averaged brain template.^44^ The diffusion sampling with a length ratio of 1.25 was used. The voxel size of the diffeomorphic reconstruction output was 2 mm isotropic. The color-coded fractional anisotropy (FA), mean diffusivity (MD), and axis-specific diffusivity (Dxx, Dyy, and Dzz) maps were estimated based on the tensor model. The DWIs exhibiting a low signal-to-noise ratio, poor spatial correlation with neighboring samples, failure in image correction, or visible artifacts were excluded, and the reconstructed images with low spatial consistency relative to the template were also removed.

### 2.4 Quantification of Periventricular Diffusivity

#### 2.4.1 DTI-ALPS calculation

To define the ROI for DTI-ALPS calculation,^25, 27^ we first created a group-averaged template in the MNI space by using the reconstructed DWIs from the HCP-Aging project^45^ accessed through the DSI studio data sharing portal (https://brain.labsolver.org/hcp_a.html). We identified projection and association fibers on the color-coded FA maps of the template and placed spherical ROIs with a 3 mm radius in these fiber areas, specifically at the level of the lateral ventricle body on both sides of the brain (Supplementary Material S2). A total 4 ROIs were applied to the axis-specific diffusivity maps of each subject, and the diffusivity values of Dxx, Dyy and Dzz within those ROIs were sampled for the DTI-ALPS calculation. The DTI-ALPS is a ratio of the average axis-specific diffusivities in the area of projection fibers (Dxx−proj) and association fibers (Dxx−assoc) on the x-axis to that of the projection fibers (Dyy−proj) on the y-axis and the association fibers (Dzz−assoc) on the z-axis (Supplementary Material S2). The DTI-ALPS indices (left-side, right-side, and mean values) were calculated for each subject to reflect the diffusion signals related to the glymphatic system.^25^

#### 2.4.2 Estimation of PVeD

Inspired by the DTI-ALPS calculation,^25^ we devised an automatic method to approximate the transverse portion of fluid diffusivity in the periventricular area (i.e. PVeD) that is hypothesized to reflect fast diffusion process in the perivenous space (Figure 2), assessing whether the derived measure could reflect biosignals related to the clearance integrity of the glymphatic system.^46^ To capture the fast diffusion signals related to the ISF dynamics along with the perivenous space around deep medullary veins that are predominantly oriented in the left-right direction, we defined an image contrast called transverse tensor ratio (TTR), which is a voxel-wise ratio calculated by dividing the tensor element on the x-axis (Dxx) by the Euclidean norm of tensor elements on three axes (Dxx, Dyy, and Dzz), capturing the transverse portion of water diffusivity on a voxel basis compared to other orthogonal directions. We also developed an automatic region growing algorithm to label periventricular regions based on the MD maps to target the periventricular areas that contain deep medullary veins. In practice, the MD maps were segmented to generate white matter (WM) masks, and a predefined lateral ventricle (LV) prior mask would be registered onto the MD maps to contour individual LV morphology, and the generated LV posterior mask would be used to create initial periventricular areas (PVeAs) as the sampling ROI through an optimized region growing algorithm. The algorithm implemented an anatomically-guided dilation of the lateral ventricle (LV) posterior mask along its transverse axis, with differential expansion coefficients that correspond to the known lateral ventricular morphology. Maximum dilation occurs at the ventricular body, while minimal expansion is applied at the peripheral ventricular margins. The hyperparameters of the region growing algorithm were empirically optimized through a multimodal image analysis, which incorporated susceptibility-weighted images, T1-weighted images, and DWIs to achieve optimal delineation of the periventricular region with specific emphasis on deep medullary vein localization (Supplementary Material S3). The initial PVeA mask was further overlapped with the WM mask to ensure that the ROI only contains the WM parenchyma, and their intersection was the final PVeA mask, which was used to sample the TTR values. Here, we calculated the median value of the TTR within the final PVeA mask to represent the average transverse portion of interstitial fluid diffusivity in the PVeA, namely periventricular diffusivity (PVeD). Processing was performed using the SPM12 and CAT12 packages ^37, 47^ in MATLAB (R2023a) (MathWorks, Natick, MA, USA). The scripts and codes of PVeD estimation are online available through our GitHub repository (https://github.com/ChangleChen/EstPVeD).

**Figure 2.**
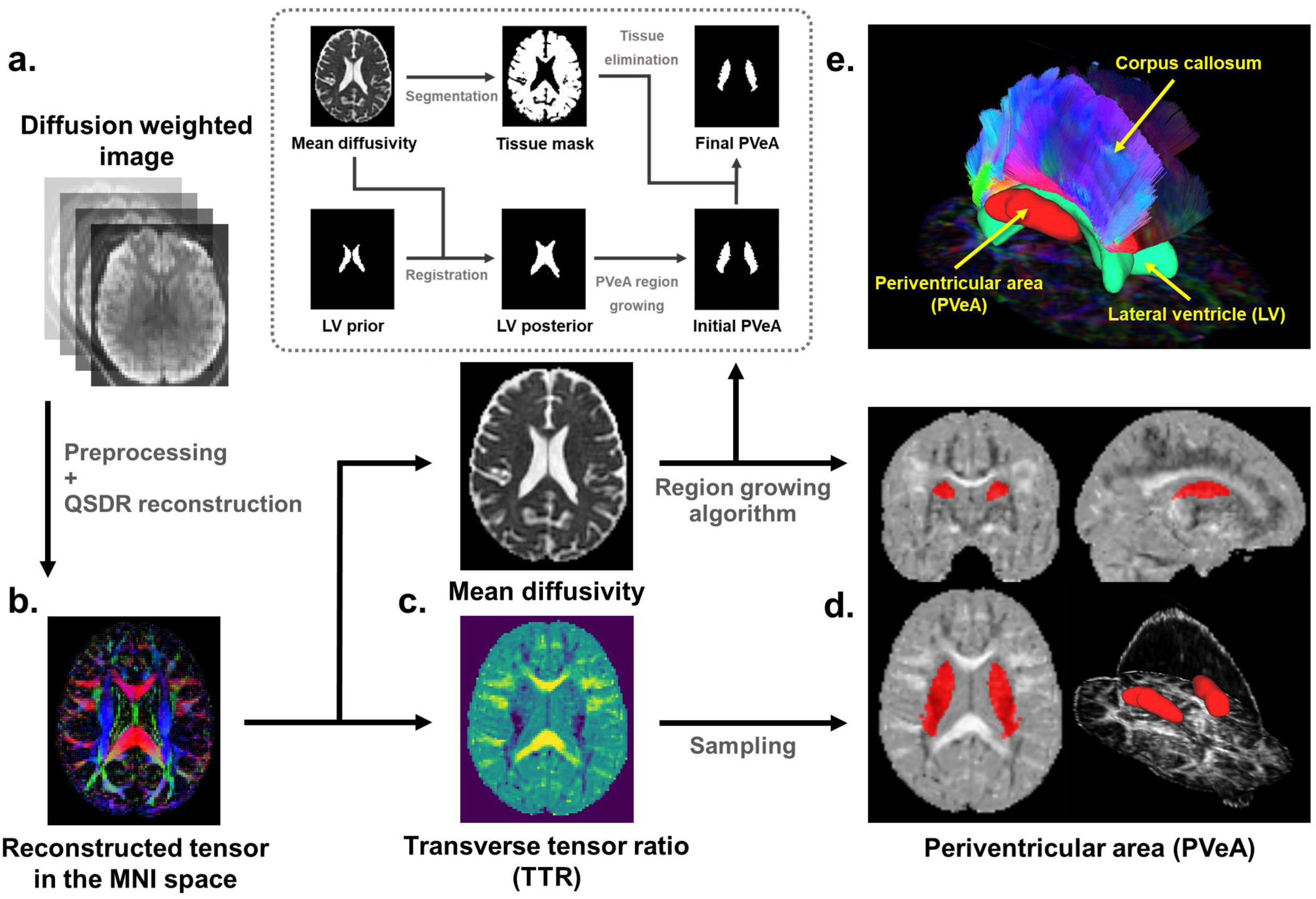
Flowchart illustrating the proposed method for estimating the transverse portion of periventricular diffusivity (PVeD) Diffusion-weighted images (DWIs) were preprocessed including phase distortion, eddy current, and motion corrections (a). The corrected DWIs were reconstructed using q-space diffeomorphic reconstruction (QSDR) to map images into the MNI space and estimate diffusion tensors (b). Meanwhile, mean diffusivity (MD) and axis-specific diffusivity maps (Dxx, Dyy, and Dzz) were calculated (c). By using axis-specific diffusivity maps, a novel transverse tensor ratio (TTR) contrast was defined to reflect transverse water diffusivity potentially along the perivascular space in deep medullary veins (c). Periventricular area (PVeA) was identified using an automated region growing algorithm applied to MD maps, incorporating lateral ventricle priors and anatomically guided dilation (d). The final PVeA mask was intersected with white matter masks, and average TTR values within the PVeA mask were calculated to approximate the apparent PVeD (e), which is used as an overall metric to represent interstitial fluid properties that could be related to glymphatic function.

### 2.5 Neuropsychological, Laboratory, and Clinical Assessments

#### 2.5.1 Neuropsychological measurement

For the BICWALZS cohort, cognitive assessment was conducted using a comprehensive battery of standardized instruments. General cognitive status was evaluated using the MMSE.^48^ Dementia severity was assessed using the Clinical Dementia Rating Sum of Boxes (CDR-SB) and the Global CDR score.^49^ Domain-specific cognitive functions were evaluated using the Seoul Neuropsychological Screening Battery,^50^ which comprises validated neuropsychological measures across multiple cognitive domains. Language function was examined using the Korean version of the Boston Naming Test (K-BNT). Visuospatial abilities and memory were assessed using the Rey Complex Figure Test and Recognition Trial (RCFT). Verbal memory was measured using the Seoul Verbal Learning Test Elderly’s version (SVLT-E). Executive function, specifically inhibitory control, was evaluated using the Korean version of the Color Word Stroop Test Color Reading (K-CWST CR). All cognitive test scores were standardized to z-scores using demographic-matched healthy controls as the reference. A subset of the participants had the second time-point measures of MMSE, which would be used to correlate with the imaging-derived measures.

Participants in the MCSA were evaluated by a comprehensive neuropsychological battery comprising 9 standardized tests to assess 4 cognitive domains^34^: (1) memory, evaluated using the Auditory Verbal Learning Test Delayed Recall Trial as well as the Logical Memory II and Visual Reproduction II subtests from the Wechsler Memory Scale-Revised (WMS-R); (2) language, assessed with the Boston Naming Test and the Category Fluency Test; (3) attention/executive function, measured using the Trail Making Test Part B and the Digit Symbol subtest of the Wechsler Adult Intelligence Scale-Revised (WAIS-R); and (4) visuospatial abilities, assessed through the Picture Completion and Block Design subtests of the WAIS-R. Raw scores for each test were standardized into z-scores, which were then averaged within each domain to compute domain-specific cognitive z-scores (i.e. memory, language, attention/executive function, and visuospatial abilities). A global cognitive z-score, reflecting overall cognitive performance, was derived by averaging the 4 domain-specific z-scores. Similarly, general cognitive status was evaluated using the MMSE, and dementia severity was assessed using the CDR-SB and Global CDR scores. We used the domain-specific cognitive scores, MMSE, Global CDR, and CDR-SB (variable name *pzmemory, pzlanguage, pzattention, pzvisualsp, MMSEcalc, CDRGLOB, and CDRSUM*) provided by the MCSA online data repository for analyses. The participants also underwent longitudinal assessment of MMSE, which was utilized to examine correlations with imaging-derived measures.

#### 2.5.2 APOE genotyping

All study participants in the BICWALZS provided written informed consent for blood collection and genomic DNA analysis. Genomic DNA was extracted from peripheral blood samples, and single-nucleotide polymorphism (SNP) genotyping was conducted using the Affymetrix Axiom KORV1.0-96 Array (Thermo Fisher Scientific, Waltham, MA, USA). All genotyping procedures were performed by DNA Link, Inc. (Seoul, Korea) in accordance with the manufacturer’s standardized protocols. APOE genotypes were determined through the analysis of two SNPs: rs429358 and rs7412, both of which were incorporated in the genotyping array. The participants with APOE4 allele were labeled as APOE4 carriers.

In the MCSA dataset, blood samples from the participants collected in clinic were used to determine APOE genotype. We used the binary status of APOE4 allele positivity (variable name *Any_E4*) provided by the MCSA online data repository for analyses.

#### 2.5.3 Clinical assessment

T1-weighted images acquired in the BICWALZS were evaluated by trained clinicians to assess medial temporal lobe atrophy (MTA) scores. To minimize intra- and inter-rater variability, the assessments were conducted by a team of two psychiatrists and six neurologists who underwent standardized training in the visual rating procedure.^51^ Coronal T1-weighted images were utilized for the evaluation with left and right MTA scored independently. The severity of MTA was graded on a standardized scale ranging from 0 (no atrophy) to 4 (severe atrophy),^52^ which was used to represent medial temporal lobe degeneration.

### 2.6 Statistical Analysis

Since the neuroimaging data were acquired at multiple sites, to mitigate the scanner-related variability,^53^ the ComBat harmonization procedure was performed to harmonize the imaging-derived measures (e.g. DTI-ALPS, PVeD, PET CL SUVR, and hippocampal volume) across scanners, while preserving the information of biological covariates used in the statistical analyses for each dataset.^54,55^

All statistical analyses were performed by using MATLAB R2023a (The MathWorks Inc., Natick, MA, USA) and R software (version 4.4.0). Images that passed quality assurance procedures were used to generate imaging-derived data, and tabular data were screened for outliers; values exceeding three standard deviations (SDs) from the mean were excluded from analysis. Data analyzed in this study are reported in tables as mean (±SD) or number (%) unless otherwise specified. The chi-square test (for categorical variables), two-sample *t* test (for continuous variables with two groups), and one-way analysis of variance (ANOVA, for continuous variables with three group) were used to test group difference for demographic characteristics. Group comparisons of neuropsychological and imaging characteristics were examined by using analysis of covariance (ANCOVA) controlling for age, sex, and education.

To explore the association of PVeD with amyloid-beta deposition and other core AD characteristics, we conducted an exploratory partial Spearman correlation analysis in the BICWALZS dataset while adjusting age, sex, and education. The core AD characteristics covered three domains including symptom severity (CDR-SB and Global CDR), cognitive outcomes (MMSE, K-BNT, RCFT, SVLT-E, and K-CWST CR), and neurodegeneration (average hippocampal volume and MTA scores). In addition, we also performed the same analysis on DTI-ALPS metrics and conventional DTI metrics including FA and MD for comparison. Given that previous studies have suggested that MD may act as a potential confounding factor in the interpretation of DTI-ALPS,^29^ we conducted an additional correlation analysis while adjusting for MD values extracted from the ROIs used in DTI-ALPS and PVeD calculations. The significance threshold was adjusted for multiple comparisons using the Bonferroni method, accounting for the combination of image feature types (i.e. DTI-ALPS, PVeD, MD, and FA) and the number of domains (i.e. symptom severity, cognitive outcomes, neurodegeneration, and Aβ burden) to mitigate Type I error inflation in the exploratory analysis. A *post hoc* analysis was conducted to examine bilateral alterations in DTI-ALPS and PVeD measures. The partial correlation analysis was also applied to the MCSA replication cohort.

If a significant association between PVeD and Aβ burden was observed, a mediation analysis was conducted to assess whether Aβ deposition mediated the relationship between PVeD and cognitive change. The mediation model was structured as a three-variable regression framework, with PVeD as the independent variable, cognitive performance as the dependent variable, and amyloid-beta deposition as the mediator. Covariates including age, sex, and education were controlled in the analysis. Unstandardized regression coefficients were reported for each path, and the proportion of mediation (PM) was calculated to quantify the relative contribution of the indirect effect to the total effect. Multiple comparison correction was applied using the Benjamini-Hochberg method to account for the number of image measures. All analyses were performed using the *lavaan* package (version 0.6.18) in R.

To examine the effect of APOE4 status on the association between PVeD and Aβ deposition, we employed robust local weighted regression modeling in which amyloid CL SUVR values served as the dependent variable while PVeD, APOE4 positivity, and their interaction were included as independent variables. Additional covariates including age, sex, and education were also included. The Benjamini-Hochberg method was applied for multiple comparison correction given the number of image measures. Furthermore, the interaction analysis was validated in the MCSA cohort to assess the genetic modulation of APOE4 on the PVeD-Aβ relationship across the Alzheimer’s disease continuum.

To assess whether PVeD metrics at baseline could predict longitudinal cognitive change, we collected a subset of participants who had the second time-point measurement of MMSE from the BICWALZS study and calculated the annual change rate of MMSE by subtracting the baseline MMSE score from the second time-point measurement and then dividing by the time interval in years. Linear regression analysis was conducted in which the MMSE annual change rate was the dependent variable while the independent variables included baseline PVeD and other covariates involving baseline age, sex, and education. The Benjamini-Hochberg method was applied to address multiple comparisons given the number of image measures. The analysis was also performed for DTI-ALPS metrics. To validate the findings, we selected participants from the MCSA who had longitudinal MMSE assessments. Since the MCSA primarily includes cognitively unimpaired individuals, we identified a subset who exhibited cognitive decline at the second time point to replicate the experiment.

## 3 Results

### 3.1 Clinical Characteristics of Study Cohorts

Table 1 summarizes the clinical characteristics of the study cohort from the BICWALZS project. Among 440 participants, 302 (68.6%) were female, and 124 (28.2%) were APOE4 carriers. The mean (±SD) of age and education were 73.1 (±6.47) and 7.78 (±4.79) years, respectively. The majority of participants were diagnosed with mild cognitive impairment (n = 275, 62.5%). The dementia group contains the participants diagnosed with Alzheimer’s disease (n = 81, 18.4%), vascular dementia (n = 28, 6.4%), frontotemporal lobe dementia (n = 4, 0.9%), and other degenerative conditions (n = 7, 1.6%). The symptom severity, neuropsychological measures, and imaging-derived phenotypes are also displayed in Table 1. Within this cohort, there were 77 participants who had longitudinal measurement of MMSE and their demographic information is shown as follows; age at baseline: 72.2 (±5.8) years, 56 females (72.7%), and education: 7.6 (±5.1) years. The MMSE scores at baseline and at the second time point were 24.1 (±4.4) and 23.8 (±5.7), respectively. The time interval was 2.0 (±0.4) years, and the average annual change rate was −0.064 (±2.15).

**Table 1.**
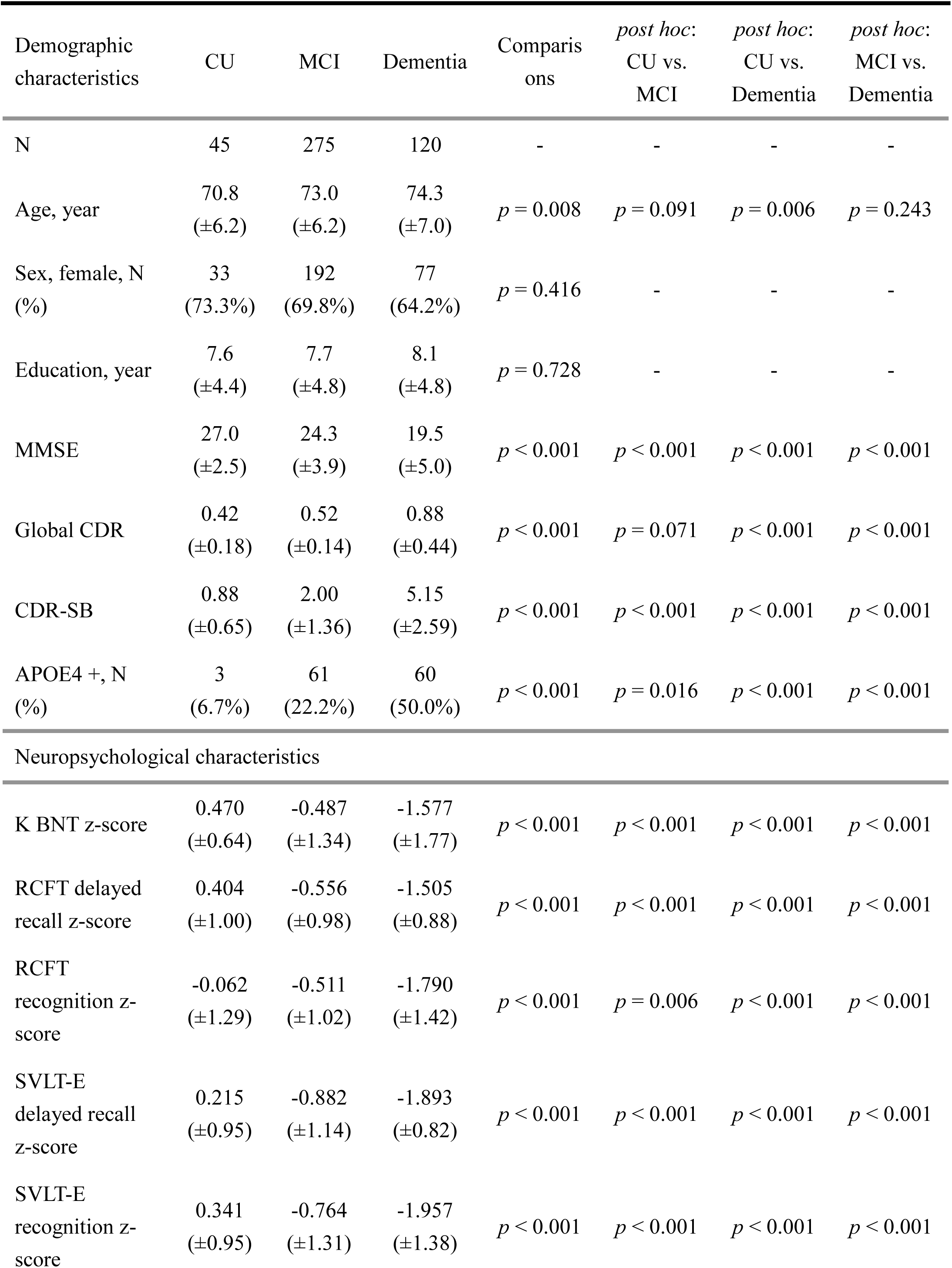
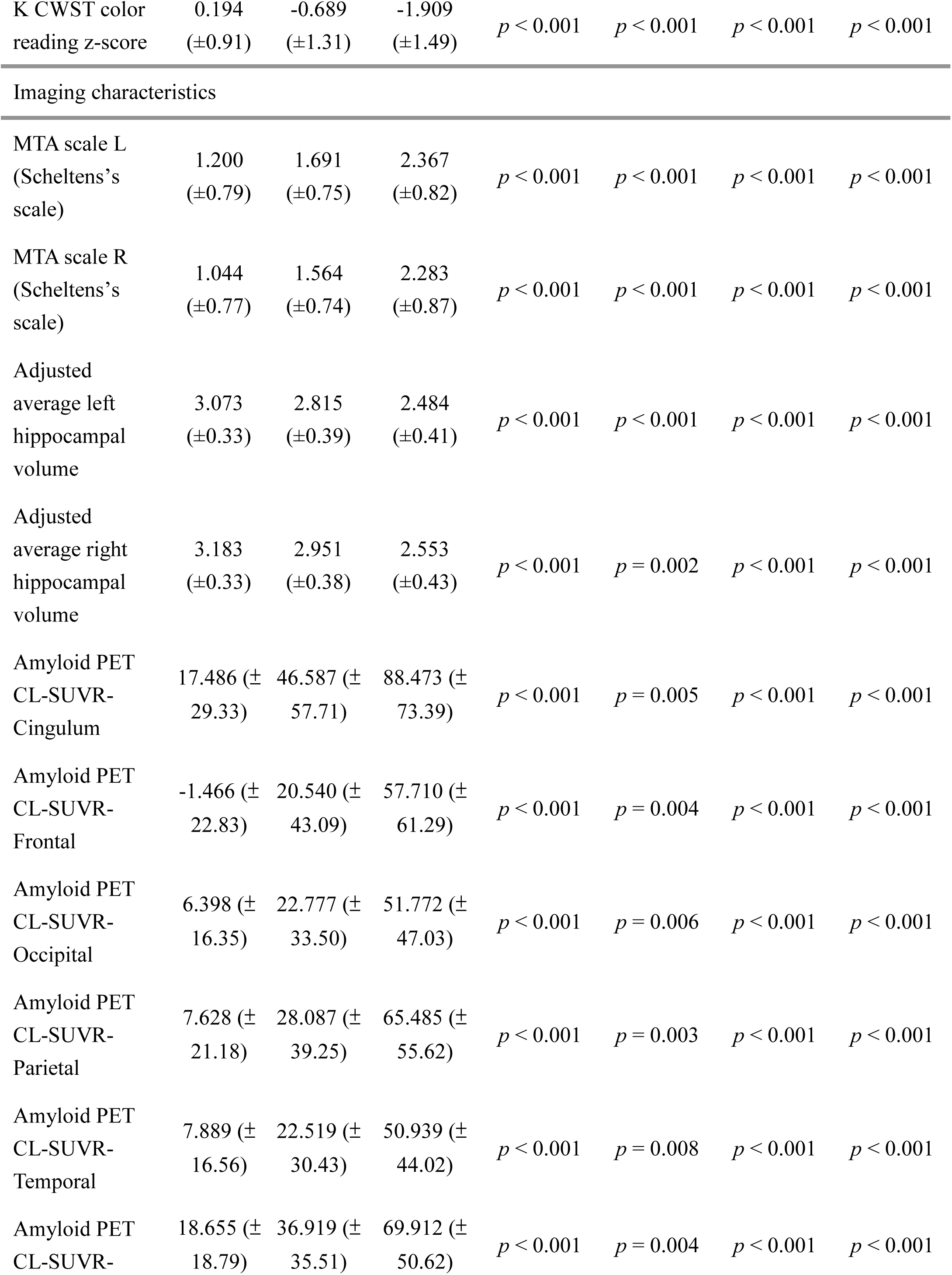
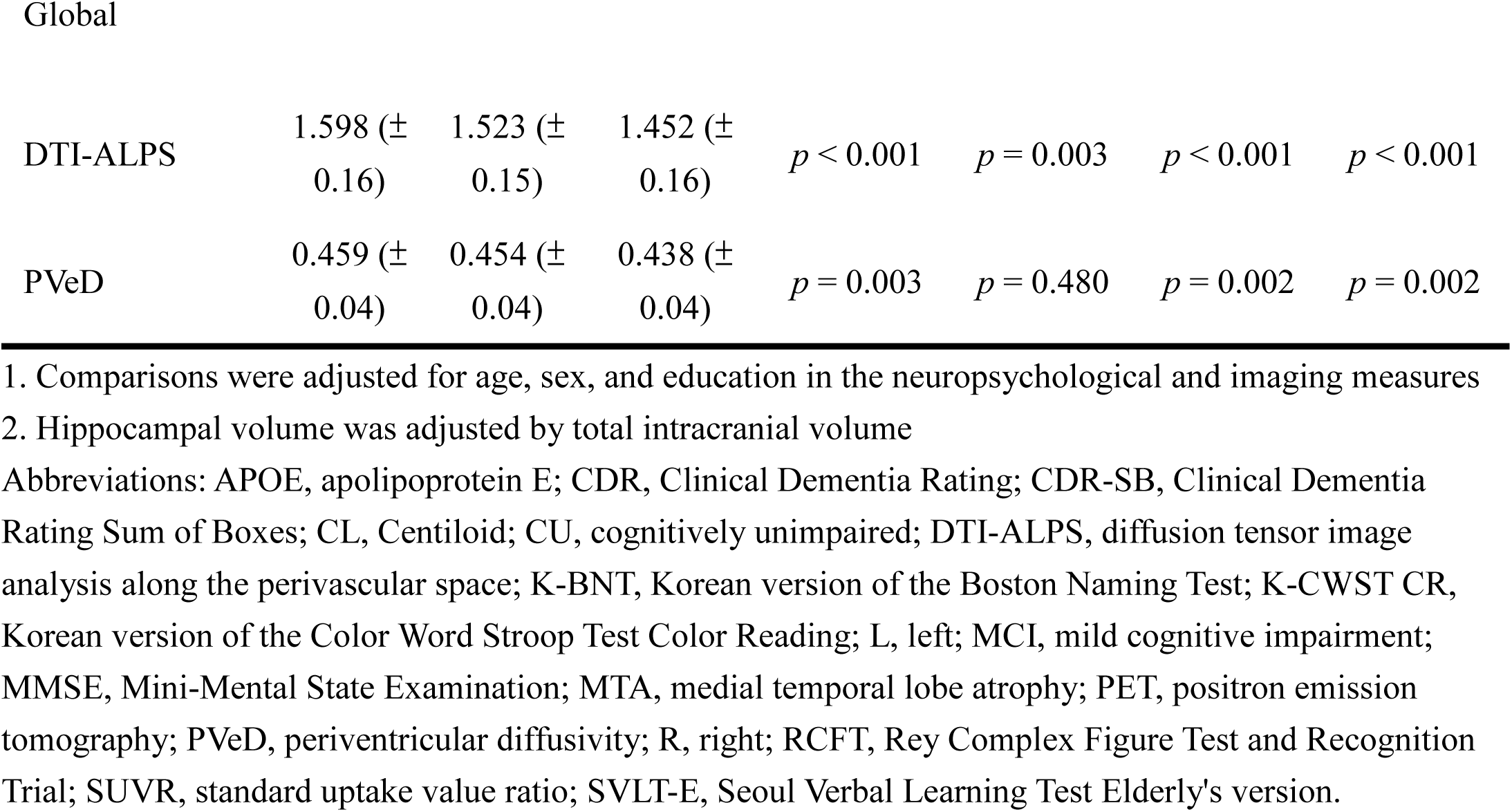
Characteristics of the study cohort from BICWALZS.

For replication, following the data quality assurance process, a total of 414 participants were selected from the MCSA study in which 351 cognitively unimpaired individuals, 57 individuals with MCI, and 6 patients diagnosed with dementia were included. Due to the limited number of participants in the dementia group, individuals with MCI or dementia were combined into one group for analysis. Among these participants, 189 (45.7%) were female, and 106 (25.6%) were APOE4 carriers. The mean (±SD) of age and education were 75.3 (±8.50) and 14.4 (±2.77) years, respectively. The details of demographics, symptom severity, neuropsychological measures, and imaging-derived phenotypes are provided in Supplementary Table 2.

### 3.2 Association of PVeD with Aβ Deposition, Neurodegeneration, Cognitive Outcomes, and Symptom Severity in the BICWALZS Cohort

The partial correlation analysis revealed that PVeD metrics were primarily correlated with symptom severity, cognitive performance, neurodegeneration, and Aβ burden after multiple comparison correction (Figure 3a & Supplementary Table 3). Lower PVeD values were associated with higher Global CDR (*P* = 2.9×10^−4^), CDR-SB (*P* = 3.8×10^−5^), MTA scores (*P* = 1.7×10^−30^ for the left side; *P* = 3.4×10^−23^ for the right side), and amyloid CL SUVRs (*P* = 8.1×10^−4^ for the global region) whereas higher PVeD values were associated with better cognitive performance (*P* = 3.5×10^−4^ for MMSE) across multiple domains and reduced hippocampal volume (*P* = 3.3×10^−17^ for the left side; *P* = 3.7×10^−18^ for the right side). These findings suggest that the interpretation of PVeD aligns with that of DTI-ALPS where higher values may indicate more efficient glymphatic-related clearance. However, PVeD demonstrated overall stronger correlations with cognitive decline, neurodegeneration, and particularly Aβ deposition (Figure 3a). Notably, the PVeD metric in the left hemisphere exhibited a preferential association with Aβ deposition compared to the right hemisphere (Figure 3b). On the other hand, the DTI-ALPS metrics demonstrated a stronger association with symptom severity and cognitive outcomes in the left-sided region compared to the right (Figure 3b). Additionally, we examined conventional DTI markers including FA and MD to show the association of white matter integrity with clinical factors (Figure 3a). Given prior reports that DTI-ALPS may be influenced by white matter diffusivity, we repeated the analysis with MD included as an additional covariate. The results indicated that the correlation patterns between PVeD and clinical factors remained robust; however, the associations between DTI-ALPS and clinical measures were diminished particularly for cognitive outcomes (Supplementary Material S4).

**Figure 3.**
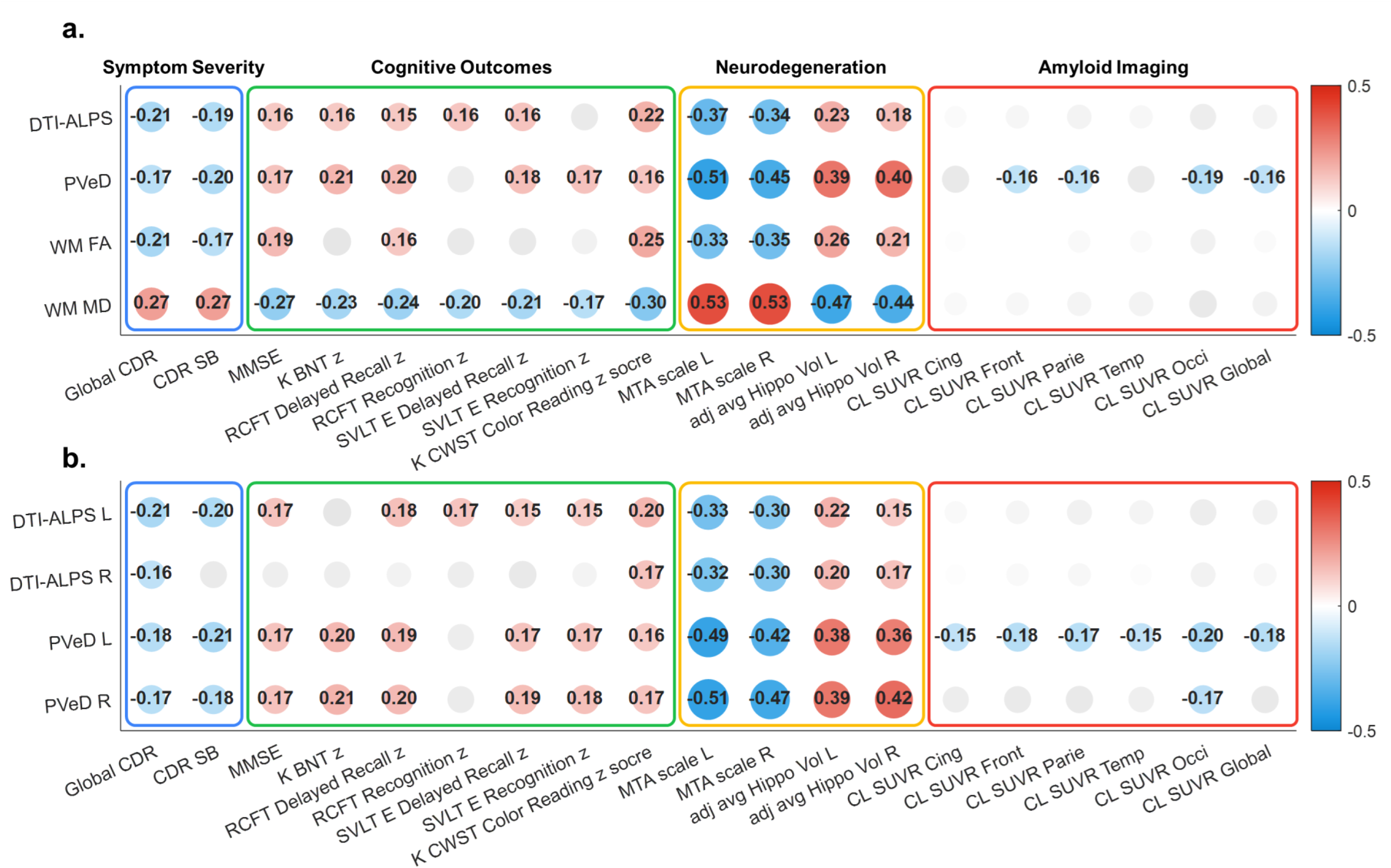
Partial correlation matrix of diffusion imaging-derived metrics with multifaceted clinical characteristics. Diffusion imaging-derived metrics refer to the diffusion tensor imaging (DTI)-derived measures including DTI-ALPS, periventricular diffusivity (PVeD), fractional anisotropy (FA), and mean diffusivity (MD). The multifaceted clinical characteristics include four domains; symptom severity, cognitive outcomes, neurodegeneration, and Aβ burden. The matrix was generated using the BICWALZS dataset (n = 440). Panels (a) and (b) present the correlation matrix of overall diffusion imaging-derived metrics and that of bilateral metrics for DTI-ALPS and PVeD, respectively. Values are Spearman’s correlation coefficients for each pairwise correlation. Covariates including age, sex, and education were adjusted. Hippocampal volume was additionally adjusted by the total intracranial volume. Darker blue and red colors indicate greater negative and positive correlations, respectively. Gray dots indicate those did not pass the significance threshold adjusted by multiple comparisons (adjusted alpha threshold = 0.0031). Abbreviations: adj, adjusted; avg, average; CDR, Clinical Dementia Rating; CDR-SB, Clinical Dementia Rating Sum of Boxes; Cing, cingulum; CL, Centiloid; CU, cognitively unimpaired; DTI, diffusion tensor image; DTI-ALPS, diffusion tensor image analysis along the perivascular space; Front, frontal; Hippo, hippocampal; K-BNT, Korean version of the Boston Naming Test; K-CWST, Korean version of the Color Word Stroop Test; L, left; MCI, mild cognitive impairment; MMSE, Mini-Mental State Examination; MTA, medial temporal lobe atrophy; Occi, occipital; Parie, parietal; PET, positron emission tomography; PVeD, periventricular diffusivity; R, right; RCFT, Rey Complex Figure Test and Recognition Trial; SUVR, standard uptake value ratio; SVLT-E, Seoul Verbal Learning Test Elderly’s version; Temp, temporal; Vol, volume; WM, white matter; z, z-score.

### 3.3 Mediation of Amyloid Accumulation between PVeD and Cognition in the BICWALZS Cohort

Given the observed significant associations of PVeD with cognitive outcomes and Aβ deposition, we further investigated whether amyloid accumulation could potentially serve as a mediator in the relationship between PVeD and cognitive decline. The mediation analysis in the BICWALZS cohort demonstrated that both PVeD and DTI-ALPS significantly and directly explained cognitive changes represented by MMSE and CDR-SB (Figure 4). However, global Aβ burden represented by global PET CL SUVR partially mediated the relationship only between PVeD and cognitive decline (*P*_FDR_ = 0.028 for MMSE, Figure 4a; *P*_FDR_ = 0.028 for CDR-SB, Figure 4b) whereas no significant mediation effect was observed in the relationship between DTI-ALPS and cognitive decline (*P*_FDR_ = 0.196 for MMSE, Figure 4c; *P*_FDR_ = 0.196 for CDR-SB, Figure 4d). These findings indicate that lower PVeD is associated with lower MMSE scores and higher CDR-SB scores where this relationship is partially mediated by greater global Aβ deposition. While DTI-ALPS exhibited a similar pattern of cascade correlations, only the direct effect remained significant, which was likely due to the lack of a significant association between DTI-ALPS and global PET CL SUVR (*P*_FDR_ = 0.182).

**Figure 4.**
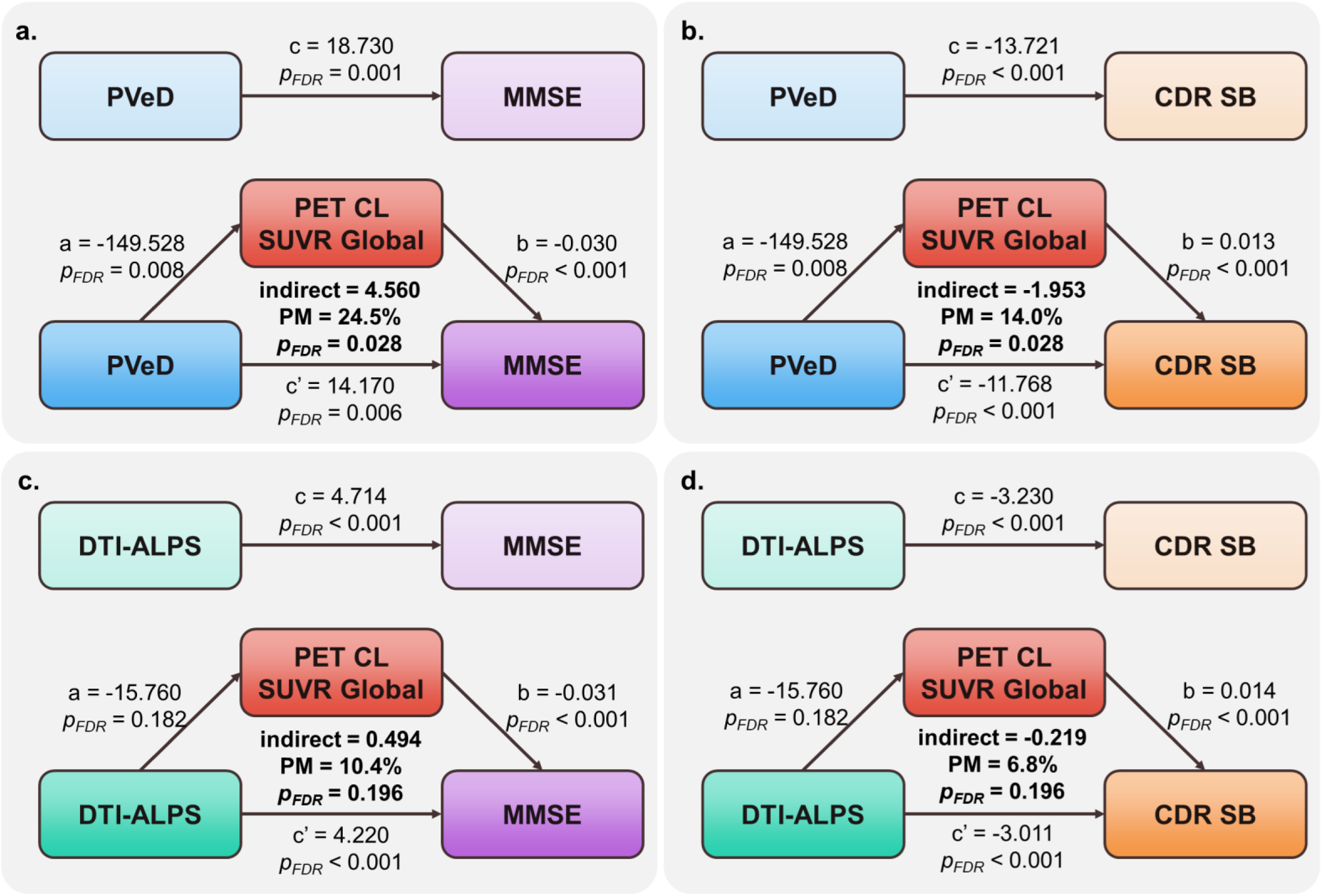
Aβ burden mediates the relationship between periventricular diffusivity (PVeD) and cognitive decline in the BICWALZS cohort. Aβ burden is quantified using the global amyloid PET standard uptake value ratio (SUVR), and cognitive decline is represented by Mini-Mental State Examination (MMSE) scores and Clinical Dementia Rating Sum of Boxes (CDR-SB) scores. Panels (a), (b), (c), and (d) illustrate the conditions of the mediation analysis for the following variable pairs: PVeD vs. MMSE, PVeD vs. CDR-SB, DTI-ALPS vs. MMSE, and DTI-ALPS vs. CDR-SB, respectively. Unstandardized coefficients are reported, and the proportion of mediation (PM) represents the indirect effect relative to the total effect. Multiple comparisons are controlled using the Benjamini-Hochberg correction accounting for the number of imaging features tested. Abbreviations: CDR-SB, Clinical Dementia Rating Sum of Boxes; CL, Centiloid; DTI-ALPS, diffusion tensor image analysis along the perivascular space; FDR, false discovery rate; MMSE, Mini-Mental State Examination; PET, positron emission tomography; PM, proportion of mediation; PVeD, periventricular diffusivity; SUVR, standard uptake value ratio.

### 3.4 APOE4 Allele Moderates the Association between PVeD and Aβ Deposition in the BICWALZS Cohort

We further examined whether APOE4 status moderates the relationship between PVeD and Aβ deposition. We observed significant interaction effects between PVeD and APOE4 status on amyloid PET CL SUVRs across all ROIs including the cingulum (*P*_FDR_ = 5.7×10^−4^, Figure 5a), frontal lobe (*P*_FDR_ = 2.6×10^−4^, Figure 5b), parietal lobe (*P*_FDR_ = 1.3×10^−4^, Figure 5c), occipital lobe (*P*_FDR_ = 1.3×10^−4^, Figure 5d), temporal lobe (*P*_FDR_ = 1.3×10^−4^, Figure 5e), and global region (*P*_FDR_ = 1.3×10^−4^, Figure 5f). This result indicates that individuals carrying the APOE4 allele exhibit a stronger negative correlation between PVeD and Aβ burden compared to non-carriers (Figure 5 & Supplementary Table 4). In other words, for each unit decrease in PVeD, individuals carrying the APOE4 allele exhibit an additional 365.5-unit increase in global PET CL SUVR, denoting a stronger association between reduced diffusivity and higher Aβ burden in APOE4 carriers.

**Figure 5.**
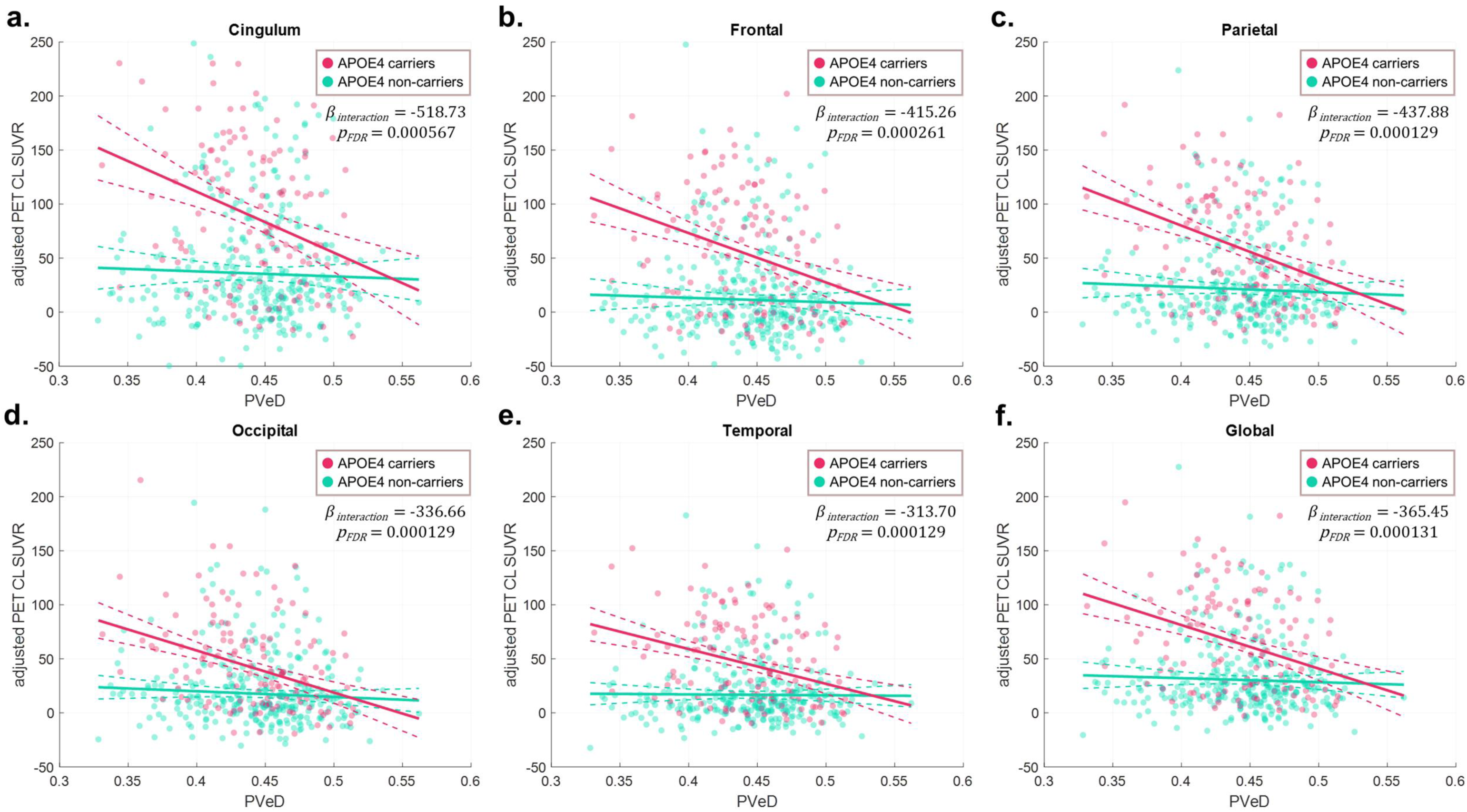
The presence of APOE4 allele enhances the association between periventricular diffusivity (PVeD) and Aβ deposition in the BICWALZS cohort. The amyloid PET CL SUVRs were sampled from five anatomical regions including the cingulum (a), frontal lobe (b), parietal lobe (c), occipital lobe (d), and temporal lobe (e), with the inclusion of the global measure (f). The models included age, sex, and education as covariates. Multiple comparisons were controlled using the Benjamini-Hochberg correction accounting for the number of image measures analyzed. Green and red colors represent APOE4 non-carriers and APOE4 carriers, respectively. The interaction terms between PVeD and APOE4 are reported. Abbreviations: APOE, apolipoprotein E; CL, Centiloid; FDR, false discovery rate; PET, positron emission tomography; PVeD, periventricular diffusivity; SUVR, standard uptake value ratio.

Given that both PVeD and DTI-ALPS reflect diffusion signals associated with glymphatic clearance efficiency, we repeated the analysis using DTI-ALPS. Although no significant correlation was observed between DTI-ALPS and Aβ deposition in the BICWALZS cohort, significant interaction effects between DTI-ALPS and APOE4 status on amyloid PET CL SUVRs were still present (Supplementary Material S5 & Supplementary Table 5). Notably, the interaction effects were more pronounced for PVeD compared to DTI-ALPS, suggesting that PVeD may be more sensitive in capturing the APOE4 modulation on the relationship between diffusion process and Aβ burden.

### 3.5 PVeD Predicts MMSE Change in the BICWALZS Cohort

We analyzed a subset of the BICWALZS cohort which had a second time-point MMSE measurement to assess whether baseline PVeD could predict the MMSE annual change rate. We observed that baseline PVeD metrics significantly predicted the MMSE annual change rate after adjusting for covariates (*P*_FDR_ = 7.9×10^−3^ for left PVeD, Figure 6a; *P*_FDR_ = 7.9×10^−3^ for right PVeD, Figure 6b; *P*_FDR_ = 7.9×10^−3^ for mean PVeD, Figure 6c) (Supplementary Table 6). Lower baseline PVeD values were associated with greater declines in MMSE, suggesting that PVeD may serve as a predictive biomarker for cognitive decline. We also repeated the analysis using DTI-ALPS metrics. Unlike PVeD, baseline DTI-ALPS metrics were not significantly associated with the MMSE annual change rate in this cohort (*P*_FDR_ = 5.9×10^−1^ for left DTI-ALPS, Figure 6d; *P*_FDR_ = 2.1×10^−1^ for right DTI-ALPS, Figure 6e; *P*_FDR_ = 2.6×10^−1^ for mean DTI-ALPS, Figure 6f). However, the trend in regression slopes suggested a similar interpretation where lower DTI-ALPS values appeared to correspond with greater cognitive decline, albeit without statistical significance (Figure 6).

**Figure 6.**
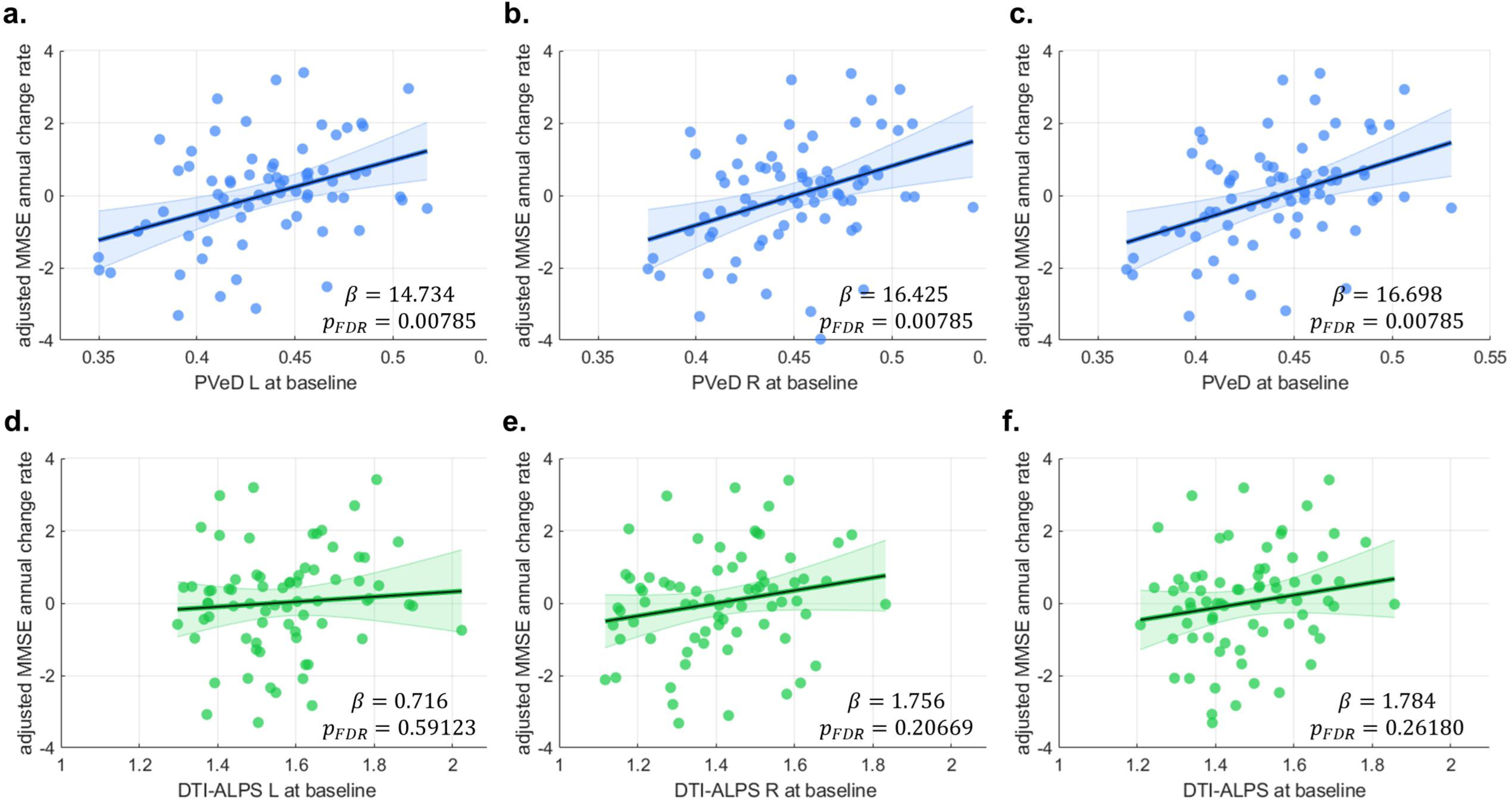
Higher baseline periventricular diffusivity (PVeD) is associated with better longitudinal cognitive outcomes in the BICWALZS cohort. The annual change rate of MMSE is used as the dependent variable with the baseline PVeDs including left PVeD (a), right PVeD (b), and mean PVeD (c) as the primary independent variable. Covariates include baseline age, sex, and education. The Benjamini-Hochberg correction was applied to account for multiple comparisons given the number of imaging measures analyzed. For reference, the analysis was also performed using DTI-ALPS metrics including left DTI-ALPS (d), right DTI-ALPS (e), and mean DTI-ALPS (f). The coefficients of slope are reported. Abbreviations: DTI-ALPS, diffusion tensor image analysis along the perivascular space; FDR, false discovery rate; MMSE, Mini-Mental State Examination.

### 3.6 Group Comparisons of PVeD and DTI-ALPS in the BICWALZS Cohort

We also conducted group comparisons of PVeD using ANCOVA while adjusting for age, sex, and education in the BICWALZS cohort (Supplementary Material S6 & Supplementary Table 7). The results demonstrated that all PVeD metrics including mean, left, and right PVeD exhibited significant differences among the groups; mean PVeD: *F*_(2,434)_ = 8.14, *P*_FDR_ = 3.8×10^−4^, left PVeD: *F*_(2,434)_ = 8.10, *P*_FDR_ = 3.8×10^−4^, and right PVeD: *F*_(2,434)_ = 8.03, *P*_FDR_ = 3.8×10^−4^. The *post hoc* analysis further showed that the significant differences existed between CU and dementia (mean PVeD: CU = 0.459 (±0.037), dementia = 0.438 (±0.039), *P_Bonferroni_* = 5.4×10^−3^; left PVeD: CU = 0.453 (±0.041), dementia = 0.430 (±0.042), *P_Bonferroni_* = 5.6×10^−3^; right PVeD: CU = 0.465 (±0.035), dementia = 0.445 (±0.040), *P_Bonferroni_* = 5.9×10^−3^) and between MCI and dementia (mean PVeD: MCI = 0.454 (±0.037), *P_Bonferroni_* = 7.0×10^−4^; left PVeD: MCI = 0.446 (±0.040), *P_Bonferroni_* = 8.0×10^−4^; right PVeD: MCI = 0.461 (±0.038), *P_Bonferroni_* = 7.0×10^−4^). There was no significant difference between CU and MCI (mean PVeD: *P_Bonferroni_* = 1.00; left PVeD: *P_Bonferroni_* = 1.00; right PVeD: *P_Bonferroni_* = 1.00). Similarly, all DTI-ALPS metrics exhibited significant differences among groups; mean PVeD: *F*_(2,434)_ = 16.12, *P*_FDR_ < 1.0×10^−6^, left PVeD: *F*_(2,434)_ = 15.62, *P*_FDR_ < 1.0×10^−6^, and right PVeD: *F*_(2,434)_ = 8.65, *P*_FDR_ = 3.8×10^−4^. The *post hoc* analysis showed that the significant differences existed between each pair of groups; (1) between CU and dementia: mean DTI-ALPS: CU = 1.598 (±0.162), dementia = 1.452 (±0.163), *P_Bonferroni_* < 1.0×10^−4^; left DTI-ALPS: CU = 1.695 (±0.197), dementia = 1.519 (±0.198), *P_Bonferroni_* = 1.0×10^−4^; right DTI-ALPS: CU = 1.520 (±0.189), dementia = 1.400 (±0.171), *P_Bonferroni_* = 1.1×10^−3^, (2) between MCI and dementia: mean DTI-ALPS: MCI = 1.523 (±0.149), *P_Bonferroni_* < 1.0×10^−4^; left DTI-ALPS: MCI = 1.606 (±0.186), *P_Bonferroni_* = 1.0×10^−4^; right DTI-ALPS: MCI = 1.457 (±0.168), *P_Bonferroni_* = 6.4×10^−3^, and (3) between CU and MCI: mean PVeD: *P_Bonferroni_* = 8.4×10^−3^; left PVeD: *P* = 1.1×10^−2^; right PVeD: *P_Bonferroni_* = 8.7× 10^−2^.

### 3.7 Replication in the MCSA Preclinical Cohort

To validate the findings identified in the BICWALZS cohort, we utilized data from the MCSA cohort to replicate the analyses. Partial correlation analysis successfully replicated the association between PVeD metrics and cognitive outcomes, confirming that lower PVeD corresponds to poorer cognitive performance (Figure 7a & Supplementary Table 8). However, the analysis did not identify significant correlations between PVeD and symptom severity, neurodegeneration, or Aβ deposition (Figure 7a). This discrepancy may be attributed to the composition of the MCSA cohort, which primarily consists of cognitively unimpaired individuals, potentially limiting the ability of PVeD to capture associations with AD signatures. A similar pattern was observed in the DTI-ALPS metrics whereas DTI-ALPS demonstrated a significant correlation with Global CDR scores (Figure 7a & Supplementary Table 8). When controlling for MD values, PVeD metrics retained their association with cognitive outcomes; however, the associations of DTI-ALPS metrics with both cognitive outcomes and symptom severity were compromised (Supplementary Figure 7a in Supplementary Material S7). Due to the absence of a significant correlation between PVeD and Aβ deposition in the MCSA cohort, mediation analysis was not conducted.

**Figure 7.**
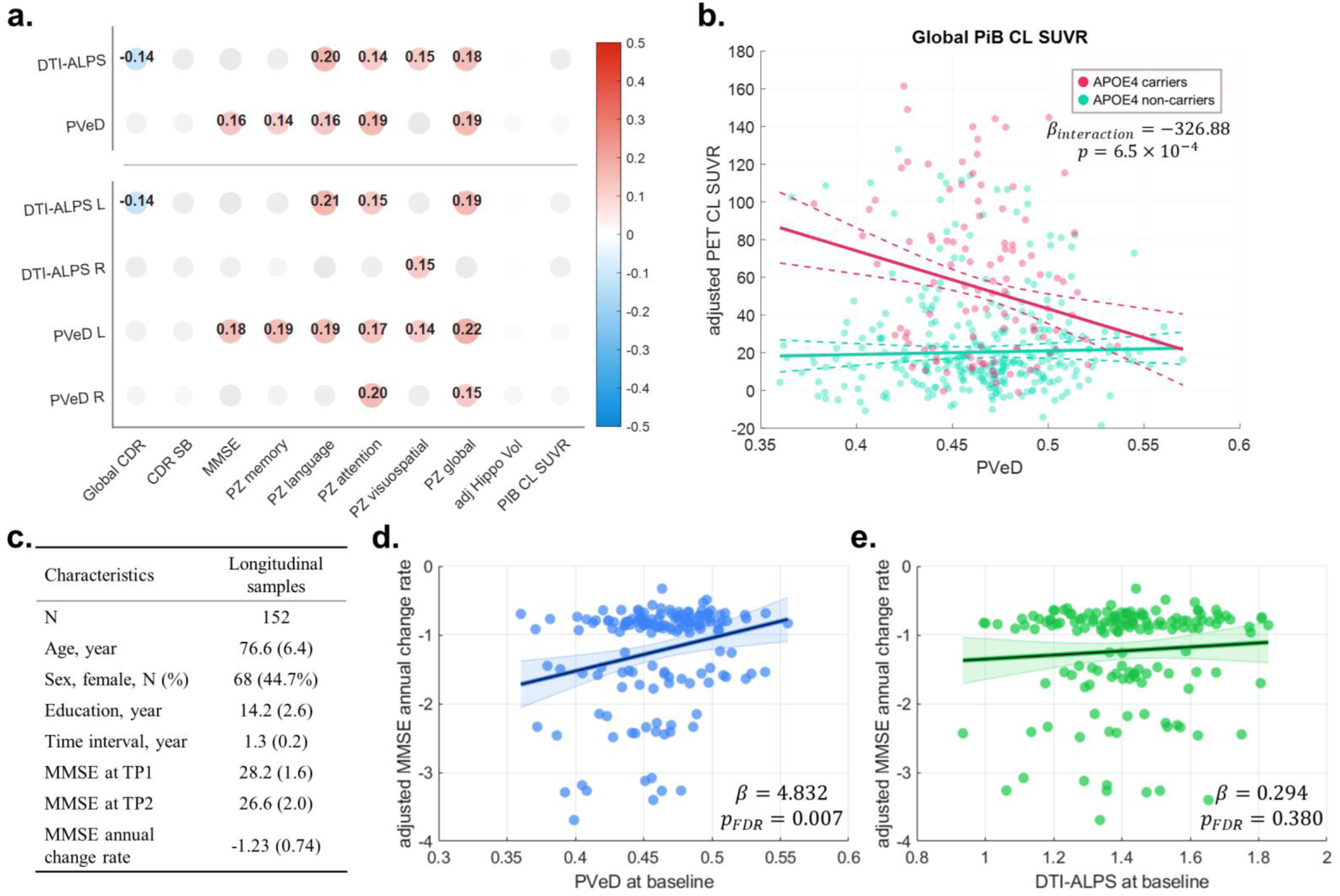
Replication of findings identified in the BICWALZS cohort using the MCSA cohort. The results including partial correlation analysis (a), interaction analysis (b), and regression analysis for predicting longitudinal cognitive decline were demonstrated (c-e). In the correlation matrix (a), values are Spearman’s correlation coefficients for each pairwise correlation. Covariates including age, sex, and education were adjusted. Hippocampal volume was additionally adjusted by the total intracranial volume. Darker blue and red colors indicate greater negative and positive correlations, respectively. Gray dots indicate those did not pass the significance threshold adjusted by multiple comparisons. For the interaction analysis (b), the amyloid PET global CL SUVR was dependent variable, with the primary independent variables including PVeD, APOE4 status, and their interaction. The model also included age, sex, and education as covariates. The demographic information is displayed for those with longitudinal cognitive decline in the MCSA cohort (c). In the regression analysis for this sample, the annual change rate of MMSE serves as the dependent variable while baseline PVeD and DTI-ALPS are included as the primary independent variables with covariates including baseline age, sex, and education (d & e). The coefficients of slope are reported. Abbreviations: adj, adjusted; APOE, apolipoprotein E; CDR, Clinical Dementia Rating; CDR-SB, Clinical Dementia Rating Sum of Boxes; CL, Centiloid; DTI-ALPS, diffusion tensor image analysis along the perivascular space; Hippo, hippocampal; L, left; MMSE, Mini-Mental State Examination; PiB, Pittsburgh Compound B; PVeD, periventricular diffusivity; R, right; SUVR, standard uptake value ratio; TP, time-point; Vol, volume; z, z-score.

Additionally, we validated whether APOE4 status moderates the relationship between PVeD and Aβ deposition in the MCSA cohort. We observed a significant interaction effect between PVeD and APOE4 status on amyloid PET CL SUVR (*P* = 6.5×10^−4^), replicating a genetic influence effect of APOE4 on the relationship between PVeD and Aβ deposition (Figure 7b & Supplementary Table 8). The interaction term between DTI-ALPS and APOE4 status also significantly explained amyloid PET CL SUVR (*P* = 1.9×10^−3^, Supplementary Figure 7b in Supplementary Material S7), further supporting the APOE4 status’ influence on the association between Aβ deposition and diffusion-based glymphatic-related metrics.

To validate whether baseline PVeD could predict longitudinal cognitive decline, we selected individuals from the MCSA cohort who exhibited a decline in MMSE over time (n = 152, Figure 7c). Regression analysis confirmed that the baseline mean PVeD metric significantly predicted the MMSE annual change rate after adjusting for covariates (*P*_FDR_ = 7.0×10^−3^) (Figure 7d & Supplementary Table 8), reinforcing the potential utility of PVeD as a predictive biomarker for cognitive deterioration. In contrast, the baseline mean DTI-ALPS metric was not significantly associated with the MMSE annual change rate in this cohort (*P*_FDR_ = 3.8 × 10^−1^) (Figure 7e). Cross-sectional group comparisons demonstrated that both PVeD and DTI-ALPS significantly differentiated CU individuals from those with cognitive impairment (including MCI and dementia) while adjusting for age, sex, and education (Supplementary Figure 8 in Supplementary Material S7); PVeD: CU = 0.474 (±0.035), MCI/dementia = 0.460 (±0.039), *F*_(1,408)_ = 7.38, *P*_FDR_ = 6.9×10^−3^; DTI-ALPS: CU = 1.474 (±0.176), MCI/dementia = 1.392 (±0.152), *F*_(1,408)_ = 11.38, *P*_FDR_ = 1.6×10^−3^.

## 4 Discussion

In this study, we investigated the utility of periventricular diffusivity (PVeD) as an imaging proxy to assess the relationship between diffusion signals that reflect fast diffusion processes within perivenous spaces in the periventricular region and Aβ deposition in the brain parenchyma. Leveraging data from two large-scale, multi-site cohorts, we demonstrated that PVeD is significantly associated with core AD characteristics in which lower PVeD values are correlated with higher Aβ deposition, reduced cognitive performance, and greater neurodegeneration across the AD continuum, suggesting that this imaging-derived phenotype could potentially serve as an imaging marker for AD. Furthermore, our study provides the first evidence of a replicable genetic modulation effect of the APOE4 allele on the relationship between PVeD and Aβ deposition. We observed that APOE4 carriers showed a stronger negative correlation between PVeD and Aβ burden, suggesting an interaction between genetic risk and glymphatic-related dysfunction in AD. We also found that baseline PVeD is a replicable predictor of longitudinal cognitive decline. Additionally, we developed an automated approach for measuring PVeD and have made it openly accessible to the scientific community, aiming to facilitate broader adoption and reproducibility in future research. Together, our findings advance the idea that periventricular ISF diffusion impairment may be associated with the progression of AD pathology and disease.

The observed stronger negative correlation between Aβ deposition and PVeD in APOE4 carriers compared to the weak or absent correlation in non-carriers suggests a critical role of APOE4 in modulating Aβ clearance dysfunction. Although several studies have demonstrated a correlation between Aβ deposition and the glymphatic-related marker, DTI-ALPS,^30, 31, 56^ they have primarily treated APOE4 as a covariate rather than investigating its potential modulatory role. As a result, none have explicitly explored how genetic variation may influence the relationship between glymphatic-related fluid movement and amyloid accumulation in humans. Our findings address this gap by providing novel evidence of modulation effect of APOE4, suggesting that its presence alters the association between PVeD and Aβ burden. That this finding is consistent across cohorts with MCI and cognitively unimpaired older adults underscores the genetic influence on glymphatic-related diffusion processes and amyloid clearance. The APOE4 allele is the strongest known genetic risk factor for AD, promoting Aβ accumulation and impeding its clearance through multiple mechanisms.^6, 57^ APOE4 accelerates early seeding of amyloid pathology by enhancing Aβ deposition and neuritic dystrophy.^58^ It also impairs Aβ clearance by reducing the size and function of meningeal lymphatic vessels.^6, 59^ Moreover, recent literature emphasizes the role of aquaporin-4 (AQP4) water channels in the glymphatic system, which facilitates the exchange of CSF and ISF in the brain.^60^ APOE4 has been linked to disrupted AQP4 polarization, potentially impairing perivascular clearance pathways and promoting amyloid retention.^61, 62^ Finally, APOE4 impairs Aβ clearance more than other isoforms, resulting in higher soluble Aβ concentrations in the brain ISF.^63^ Thus, the accumulated evidence suggests that the APOE4 allele has strong influence on glymphatic dysfunction in AD, which could exacerbate Aβ accumulation in the interstitial space.

As already discussed, the glymphatic system clears Aβ deposition through convective bulk flow and diffusion.^7, 15, 23^ The CSF enters periarterial spaces, driven by arterial pulsations, and mixes with ISF via AQP4 channels, which further facilitate the waste transportation including Aβ toward perivenous spaces.^2, 15, 64^ While bulk flow dominates perivascular spaces, diffusion also contributes to solute movement from high- to low-concentration areas.^2, 7, 20, 23^ This combination enhances Aβ removal, with fluid and solutes draining into meningeal lymphatics and other outflow routes. The relative contributions of convection and diffusion vary based on brain region, solute size, and other factors.^2, 7, 15^ Our results concur with prior studies^30, 31^ that the fast diffusion process observed through PVeD is associated with the Aβ deposition, and we further identified that this association could be moderated by the APOE4 allele status. Although the actual mechanism between impaired diffusion and Aβ deposition remains uncertain, our results support that altered diffusion may contribute to Aβ accumulation. This could potentially be due to structural changes in the vasculature, reduced permeability, or an increased concentration of metabolites within the perivenous space,^65–67^ leading to slowed or obstructed Aβ transport. Nevertheless, Aβ deposition itself may also obstruct pathways to the perivenous space, further diminishing diffusion efficiency.^7, 65^ However, we did not observe a significant association between PVeD and amyloid burden in the MCSA cohort, which mainly comprised cognitively normal preclinical participants, compared to the BICWALZS clinical cohort largely including MCI patients. This discrepancy may reflect that diffusion-related clearance becomes more pronounced with accumulating amyloid pathology and neurodegeneration.^30^ In preclinical stages, compensatory clearance mechanisms may still preserve so that PVeD alterations may emerge only with advancing disease burden and clinical symptoms.^68^ Further investigations are necessary to elucidate the fluid dynamics underlying these processes that govern Aβ clearance.

Recent studies have explored the potential of DTI-ALPS as a non-invasive imaging surrogate for assessing glymphatic-related dysfunction.^4, 18, 25^ Lower DTI-ALPS indices have been associated with increased Aβ deposition.^30, 31, 56^ However, the ROI seeding and calculation of DTI-ALPS are highly susceptible to various factors such as the confounding effect of deep white matter structure in the ROI, inter-rater variability, and image registration performance when automated methods are employed. These factors can introduce inconsistencies and increase sensitivity to potential confounders, potentially impacting the interpretation of the results.^20, 24, 29^ This is possibly reflected in our findings where the DTI-ALPS metrics demonstrated greater efficacy in distinguishing group differences, but were less likely to exhibit a specific relationship with Aβ deposition. Hence, we proposed a new method to extend the DTI-ALPS framework by automatically localizing the periventricular region encompassing the transverse perivascular space surrounding deep medullary veins. We also generalized the index calculation (i.e. transverse tensor ratio, TTR) to approximate the transverse component of fluid movement primarily along the perivenous space. Our results demonstrate that the proposed metrics effectively capture core AD signatures involving Aβ burden, neurodegeneration, and cognitive decline. Our findings in the mediation analysis also support this inference. We observed that the association between PVeD and amyloid burden appears to be primarily driven by the left hemisphere in the clinical cohort. Although glymphatic transport is generally presumed to be bilaterally symmetrical, subtle hemispheric variations in vascular and white matter geometry may result in asymmetric perivascular fluid dynamics. Further research is necessary to elucidate the mechanisms underlying the hemispheric asymmetry.

Cross-sectional cognitive performance and subsequent cognitive decline were significantly associated with PVeD metrics in two independent cohorts, which underlines its potential as an imaging marker of cognitive trajectory. In line with previous literature of which DTI-ALPS can predict cognitive changes,^30, 31^ PVeD values not only relate to current cognitive status but also predict future changes from baseline, signifying the sensitivity of fast diffusion signals in the periventricular region to shifts in cognitive function. This suggests that the investigation the biological underpinnings of altered periventricular diffusion processes could facilitate targeted interventions aimed at preserving cognitive health, particularly in vulnerable or at-risk populations.^1, 69, 70^

This study, while introducing a novel approach to measuring diffusion signals via the PVeD metric, has several limitations. First, although the PVeD metric holds promise as a non-invasive proxy for glymphatic-related integrity, it remains an indirect index, offering only an overall view of Brownian motion of water within white matter in the periventricular area. Furthermore, the spatial resolution constraints of clinical DTI data pose challenges for capturing the nuanced fluid flow in perivascular spaces. Because the PVeD metric relies on quantifying transverse diffusivity in the periventricular region, it may not sufficiently represent the complex, anisotropic nature of fluid transport within the perivascular space. Partial volume effects stemming from low spatial resolution can also introduce inaccuracies in PVeD estimates. Additionally, this method is limited to regions adjacent to the lateral ventricles and thus cannot capture the full spectrum of glymphatic functionality across the entire brain. Considering the regional heterogeneity of AD neuropathology, future studies should extend investigations to other AD-relevant regions and integrate additional disease-specific biomarkers to elucidate regional glymphatic integrity. Second, the PVeD metric rests on the assumption that water diffusion within the perivascular space is Gaussian, which may not accurately characterize true diffusion in this compartment. Similarly, the assumption of isotropic, fixed diffusivity for the free-water component may overlook important features of fluid dynamics in the narrow, tubular perivascular spaces. While we have illustrated how fast diffusion signals might reflect amyloid accumulation, the roles of other fluid mechanisms such as bulk flow in amyloid clearance warrant further exploration. Third, our longitudinal assessment was restricted to MMSE measurements from a subset of participants, limiting our ability to infer changes in amyloid clearance over time. Future research should adopt more extensive longitudinal protocols, incorporating other modes of neuroimaging, blood-based assays, and neuropsychological assessments to capture the evolving clinical progression and neuropathological processes. Fourth, although this study examined the influence of the APOE4 allele on the interaction between the diffusivity and Aβ deposition, it did not investigate other APOE variants, notably the potentially protective APOE2 allele. Such investigations may offer a more comprehensive understanding of the genetic factors influencing glymphatic function and amyloid pathology.

Altogether, by leveraging multi-national, multi-site, and multi-modal neuroimaging data spanning the Alzheimer’s spectrum, this study presents a novel approach to capture fast diffusion signal along the perivenous space in the periventricular region. We demonstrate critical evidence of its association with Aβ burden, neurodegeneration, and cognitive decline in the Alzheimer’s continuum. Our findings suggest that lower periventricular diffusivity is linked to increased Aβ deposition particularly in APOE4 carriers and could serve as an imaging predictor of longitudinal cognitive decline. These results indicate that PVeD may offer a non-invasive and clinically compatible tool for tracking Aβ clearance dysfunction in AD and evaluating therapeutic interventions. Moreover, the observed modulation by APOE4 status underscores an important connection between glymphatic-related diffusion processes and Aβ accumulation, warranting further investigation into individualized risk stratification. Future research should explore the mechanistic underpinnings of periventricular fluid transport and validate PVeD across diverse populations and imaging platforms. Additionally, expanding the application of PVeD to other neurodegenerative conditions may further elucidate its role in broader pathophysiological contexts. Integrating such diffusion imaging-derived metrics with multimodal neuroimaging and fluid biomarkers could enhance early detection and intervention strategies, ultimately contributing to a more comprehensive understanding of neurodegenerative disease mechanisms.

## 5 Acknowledgements

Funding: This work was supported by the National Institutes of Health/National Institute on Aging grant; the Normal Aging study (R01 2RF1AG025516). the SDAR study (R01AG067018), the PPG4 study (P01 AG025204), the SERA study (R01 AG085566), and the Harmonization study (R01 AG063752).

Resources: This study was conducted with biospecimens and data from the consortium of the Biobank Innovations for Chronic cerebrovascular disease With ALZheimer’s disease Study (BICWALZS), which was supported by the National Institute of Health research project, Republic of Korea (Project No. 2024-ER0505-00). The biospecimens and data used for this study were provided by the Biobank of Ajou University Hospital, a member of Korea Biobank Network. This work was supported by the National Research Foundation of Korea (NRF), funded by the Ministry of Science and ICT (RS-2019-NR040055). This research was supported by a grant of the Korea Dementia Research Project through the Korea Dementia Research Center (KDRC), funded by the Ministry of Health & Welfare and Ministry of Science and ICT, Republic of Korea (RS-2024-00339665). This research was supported by a grant of the Korea Health Technology R&D Project through the Korea Health Industry Development Institute (KHIDI), funded by the Ministry of Health & Welfare, Republic of Korea (HR21C1003, HR22C1734 and RS-2024-00406876).

The Mayo Clinic Study of Aging was supported by the NIH (U01 AG006786, P30 AG062677, R37 AG011378, R01 AG041851, R01 NS097495), the Alexander Family Alzheimer’s Disease Research Professorship of the Mayo Clinic, the Mayo Foundation for Medical Education and Research, the Liston Award, the GHR Foundation, the Schuler Foundation, and used the resources of the REP medical records linkage system, which is supported by the National Institute on Aging (NIA: AG 058738), by the Mayo Clinic Research Committee, and by fees paid annually by REP users. Dr. Devanand receives funding support from the National Institute on Aging (NIA) and the Alzheimer’s Association; serves as a scientific adviser for Acadia, GSK, Corium, Eisai; and serves on the data safety and monitoring board for BioXcel. Dr. Motter reports funding from NIA. Dr. Lee reports funding from NIA. Dr. Vassilaki has served as a consultant for F. Hoffmann-La Roche Ltd; she currently receives research funding from NIH and has equity ownership in Amgen, Johnson and Johnson, Medtronic, and Merck. Dr. Luchsinger receives research funding from NIH and receives a stipend from Wolters Kluwer as editor in chief of a journal. Dr. Knopman has received research support from the NIH and the Robert H. and Clarice Smith and Abigail Van Buren Alzheimer’s Disease Research Program of the Mayo Foundation; has served on a data safety and monitoring board for Lundbeck Pharmaceuticals and for the Dominantly Inherited Alzheimer Network (DIAN) study; and has been an investigator for clinical trials sponsored by Biogen, TauRx Pharmaceuticals, Lilly pharmaceuticals, and the Alzheimer’s Disease Treatment and Research Institute, University of Southern California.

## 7 Author’s Contribution

(1) Research Project: A. Conception, B. Organization, C. Execution; (2) Statistical Analysis: A. Design, B. Execution, C. Review and Critique; (3) Manuscript Preparation: A. Writing of the First Draft, B. Review and Critique. C.L. Chen: 1A, 1B, 1C, 2A, 2B, 2C, 3A; S.J. Son: 1B, 1C, 2A, 2C, 3B; N. Schweitzer: 2C, 3B; H. Jin: 2C, 3B; J Li: 2C, 3B; L. Wang: 2C, 3B; S. Yang: 2C, 3B; C.H. Hong: 1C, 3B; H.W. Roh: 1C, 3B; B. Park: 1C, 3B; J.W. Choi: 1C, 3B; Y.S. An: 1C, 3B; S.W. Seo: 1C, 3B; Y.H. Cho: 1C, 3B; S. Hong: 1C, 3B; Y.J. Nam: 1C, 3B; D.S. Minhas: 2C, 3B; C.M. Laymon: 2C, 3B; G.D. Stetten: 2C, 3B; D.L. Tudorascu: 2C, 3B; MCSA: 1C; H.J. Aizenstein: 1A, 1B, 1C, 2C, 3B; M. Wu: 1A, 1B, 1C, 2C, 3B.

## 8 Research data for this article

The BICWALZS datasets generated during and/or analyzed in the current study are available from the corresponding author with the approval of the BICWALZS team upon reasonable request. The MCSA dataset is open access and can be applied through their website (https://www.mayo.edu/research/centers-programs/alzheimers-disease-research-center/research-activities/mayo-clinic-study-aging/overview). Scripts of analytical methods and codes of imaging process are online available through our GitHub repository (https://github.com/ChangleChen/EstPVeD).

## 9 Ethical Approval

All procedures performed in this study involving human participants from the BICWALZS project were in accordance with the ethical standards of the Institutional Review Boards of Ajou University Hospital (AJOUIRB-SUR-2021-038) and with the 1964 Helsinki declaration and its later amendments or comparable ethical standards. Informed consent in the study was obtained from all individual participants who were recruited in the BICWALZS project. BICWALZS was registered in the Korean National Clinical Trial Registry (KCT0003391||Registration Date: 2018/07/04|| http://cris.nih.go.kr/cris/en/use_guide/cris_introduce.jsp). Similarly, all human participants provided informed consent for the MCSA study.

## Supplementary Materials

S1 Recruitment criteria of the BICWALZS dataset

S2 Region sampling for DTI-ALPS calculation

S3 Hyperparameter optimization for targeting periventricular area

S4 Partial correlation analysis of DTI-ALPS and PVeD adjusted for mean diffusivity

S5 Interaction of DTI-ALPS with APOE4 on amyloid deposition

S6 Group comparisons of DTI-ALPS and PVeD between CU, MCI, and dementia

S7 Additional results in the replication cohort: the MCSA dataset

### S1 Recruitment criteria of the BICWALZS dataset

The BICWALZS study’s primary objective is to facilitate the systematic collection and utilization of human biological specimens and real-world clinical data for research on subjective cognitive decline (SCD), mild cognitive impairment (MCI), Alzheimer’s disease (AD), and subcortical vascular dementia (SVaD) (Son et al., 2020). Clinical diagnoses were established using internationally recognized criteria: SCD was defined as self- or informant-reported cognitive decline without objective impairment in neurocognitive tasks (no less than −1.5 standard deviations in each domain and Clinical Dementia Rating (CDR) = 0) (CHOI et al., 2001); MCI was diagnosed based on a CDR score of 0.5 and the expanded Mayo Clinic criteria (Winblad et al., 2004); AD dementia was classified using the National Institute on Aging-Alzheimer’s Association (NIA-AA) Core Clinical Probable AD Dementia Criteria (McKhann et al., 2011); and SVaD was identified based on moderate-to-severe white matter hyperintensity (WMH) burden and diagnostic criteria outlined in the Diagnostic and Statistical Manual of Mental Disorders, Fifth Edition (DSM-5) (American Psychiatric Association and American Psychiatric Association, 2013). Patients with a history of neurological or systemic conditions such as territorial cerebral infarction, intracranial hemorrhage, Parkinson’s disease, heart failure, renal failure, or any condition that could confound study findings were excluded. Clinical assessments included CDR global scores and sum of boxes (CDR-SB), a validated measure of cognitive and functional impairment used in dementia clinical trials (Morris, 1993). In addition, comprehensive evaluations were conducted, including blood pressure, pulse pressure, body mass index, smoking status, amyloid positron emission tomography (PET), apolipoprotein E (APOE) genotyping, and standardized neuropsychological assessments. A subset of participants was further classified based on neuroimaging biomarkers including cortical amyloid burden and subcortical vascular burden, into cognitively unimpaired (CU), Alzheimer’s disease-related cognitive impairment (ADCI), or vascular cognitive impairment (VCI) groups. Participants with cortical amyloid pathology or infarctions unrelated to vascular dementia, such as those resulting from radiation injury, multiple sclerosis, vasculitis, or leukodystrophy, were excluded from CU and VCI classifications. The study included 1,013 participants, with subsets undergoing additional evaluations such as brain MRI (n = 817), brain amyloid PET (n = 713), single nucleotide polymorphism microarray analysis (n = 949), actigraphy measurement (n = 200), and patient-derived dermal fibroblast sampling (n = 175). A structured longitudinal follow-up protocol was established, with annual brief assessments for all participants and biannual comprehensive evaluations—including neuropsychological testing, brain MRI, and actigraphy measurements—for those exhibiting cortical amyloid burden, subcortical vascular pathology, APOE e4 positivity, or significant cognitive decline. Among the participants, 336 were followed longitudinally for cognitive decline using CDR-SB and clinical diagnoses. BICWALZS is registered in the Korean National Clinical Trial Registry (Clinical Research Information Service, identifier: KCT0003391) and was approved by the Institutional Review Board (IRB) of Ajou University Hospital (AJOUIRB-SUR-2021-038). Written informed consent was obtained from all participants and their caregivers.

### S2 Region sampling for DTI-ALPS calculation

To define the regions of interest (ROIs) for the diffusion tensor imaging–analysis along the perivascular space (DTI-ALPS) calculation (Liu et al., 2024; Taoka et al., 2017), we first generated a group-averaged template in the Montreal Neurological Institute (MNI) space (Supplementary Figure 1). This was achieved using the reconstructed diffusion-weighted images (DWIs) from the Human Connectome Project–Aging (HCP-Aging) dataset (Bookheimer et al., 2019), accessed via the DSI Studio data-sharing portal (https://brain.labsolver.org/hcp_a.html). Projection and association fibers were identified on the color-coded fractional anisotropy (FA) maps of the template, and spherical ROIs with a 3 mm radius were manually placed within these fiber regions, specifically at the level of the lateral ventricle body bilaterally. A total of four ROIs were applied to the axis-specific diffusivity maps of each subject, and the diffusivity values along the principal axes (Dxx, Dyy, and Dzz) were extracted for DTI-ALPS calculation. The DTI-ALPS index was derived as the ratio of axis-specific diffusivities in (1) projection (Dxx_proj) and association fibers (Dxx_assoc) along the x-axis to (2) projection fibers (Dyy_proj) along the y-axis and association fibers (Dzz_assoc) along the z-axis (Supplementary Figure 2). For each subject, DTI-ALPS indices were computed for the left and right hemispheres, as well as a mean value, to quantify diffusion signals associated with glymphatic function (Taoka et al., 2017). To validate the effectiveness of DTI-ALPS measures as described in previous literature (Huang et al., 2024; Li et al., 2024), we performed a regression analysis by using the BICWALZS cohort. The mean DTI-ALPS was an independent variable, and the Mini-Mental State Examination (MMSE) score was the dependent variable while controlling for age, sex, and education. The result showed a significant positive correlation between them (Supplementary Figure 1), which is in line with the previous findings.

**Supplementary Figure 1.**
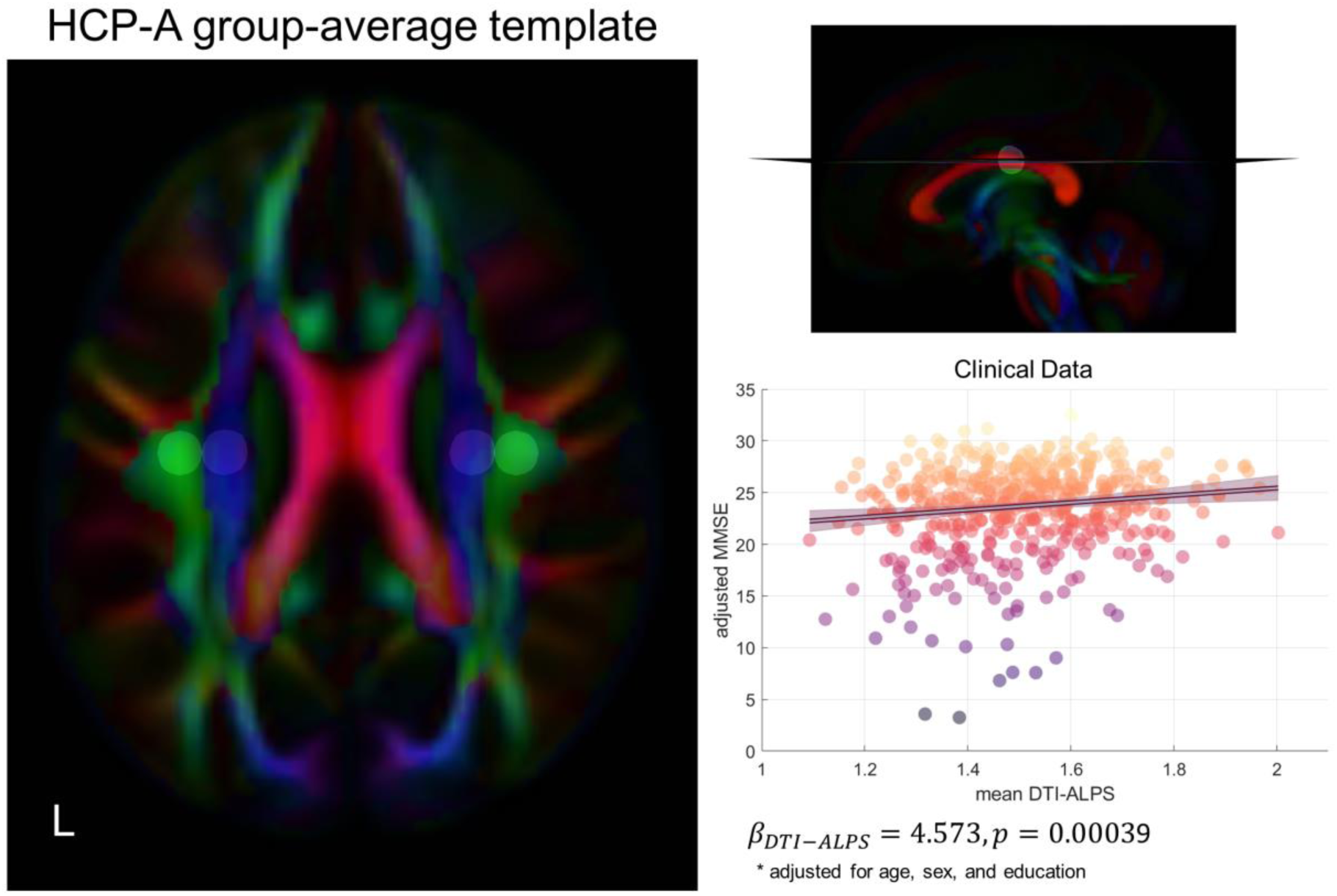
The group-average template based on the diffusion-weighted images acquired from the HCP-Aging (HCP-A) dataset for defining sampling regions of interest in the DTI-ALPS calculation.

**Supplementary Figure 2.**
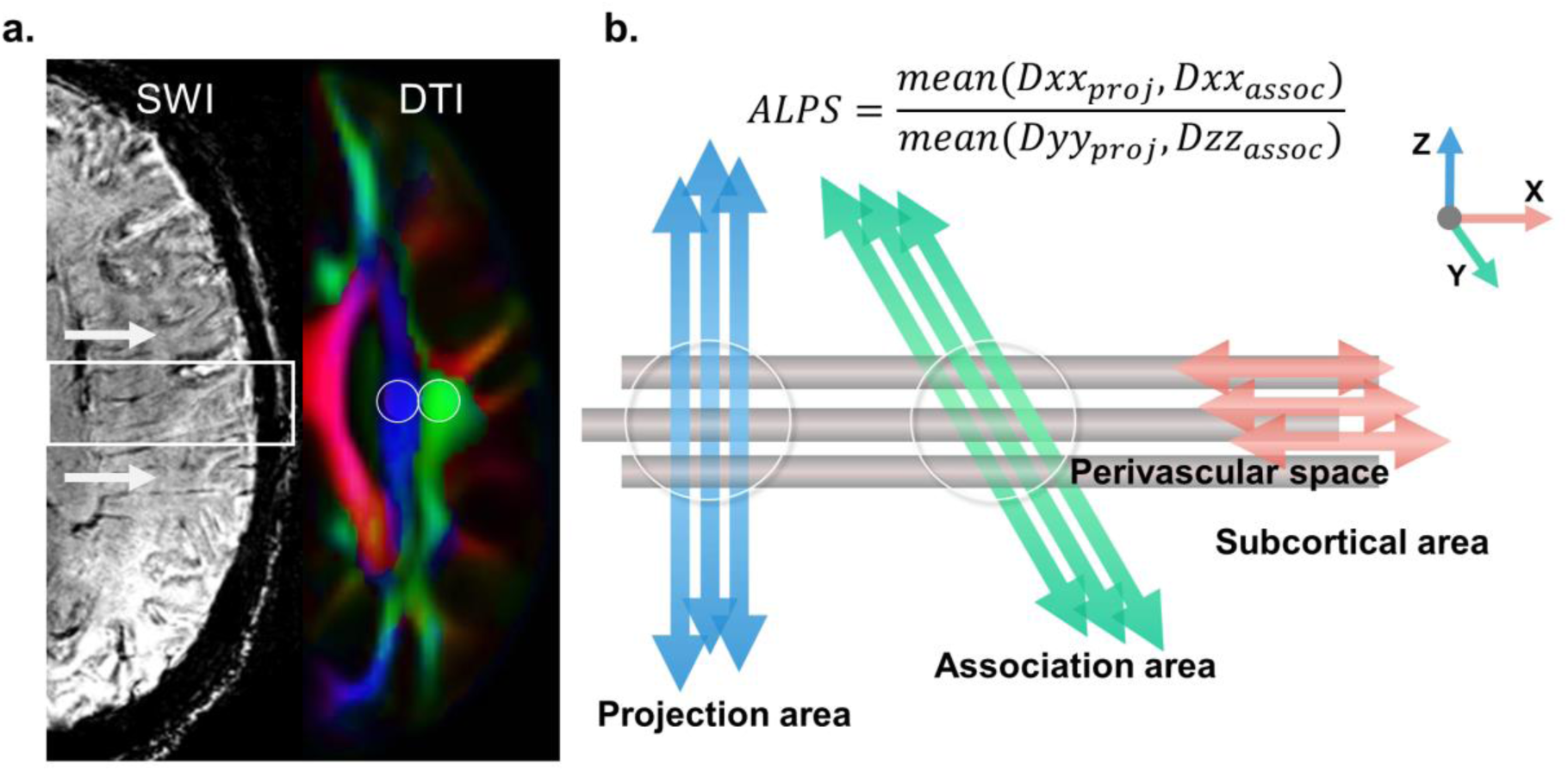
Illustration of the Diffusion Tensor Imaging Analysis along the Perivascular Space (DTI-ALPS) method designed to evaluate glymphatic function using diffusion MRI. The method quantifies diffusivity in periventricular white matter regions closely associated with the perivascular space (PVS). Spherical regions of interest are manually placed in deep white matter adjacent to the lateral ventricles (a), covering both projection and association fibers. The DTI-ALPS index is derived by comparing diffusivity along directions parallel and perpendicular to the PVS (b), providing a metric for assessing glymphatic-related function in the brain.

### S3 Hyperparameter optimization for targeting periventricular area

We developed an automated region growth algorithm to delineate periventricular regions from mean diffusivity (MD) maps, specifically targeting areas containing deep medullary veins. The algorithm first segments MD maps to generate white matter (WM) masks, followed by the registration of a predefined lateral ventricle (LV) prior mask onto the MD maps to accommodate individual LV morphology. A posterior LV mask is then derived and serves as the basis for initializing periventricular areas (PVeAs), which function as the sampling regions of interest (ROIs). To refine these ROIs, the algorithm applies an anatomically guided dilation of the LV posterior mask along the transverse axis, incorporating differential expansion coefficients that account for the known morphological variations of the lateral ventricles. Maximum dilation is applied at the ventricular body, whereas minimal expansion is enforced at the peripheral ventricular margins to preserve anatomical fidelity. The hyperparameters of the region growth algorithm were empirically optimized through multimodal image analysis, integrating susceptibility-weighted imaging, T1-weighted imaging, and diffusion-weighted imaging (DWI) to enhance the accuracy of periventricular region delineation, with particular emphasis on deep medullary vein localization (Supplementary Figure 3). This analysis was performed using a proprietary multimodal dataset, and the performance of the ROI generation process was rigorously assessed by experienced neuroimaging experts.

**Supplementary Figure 3.**
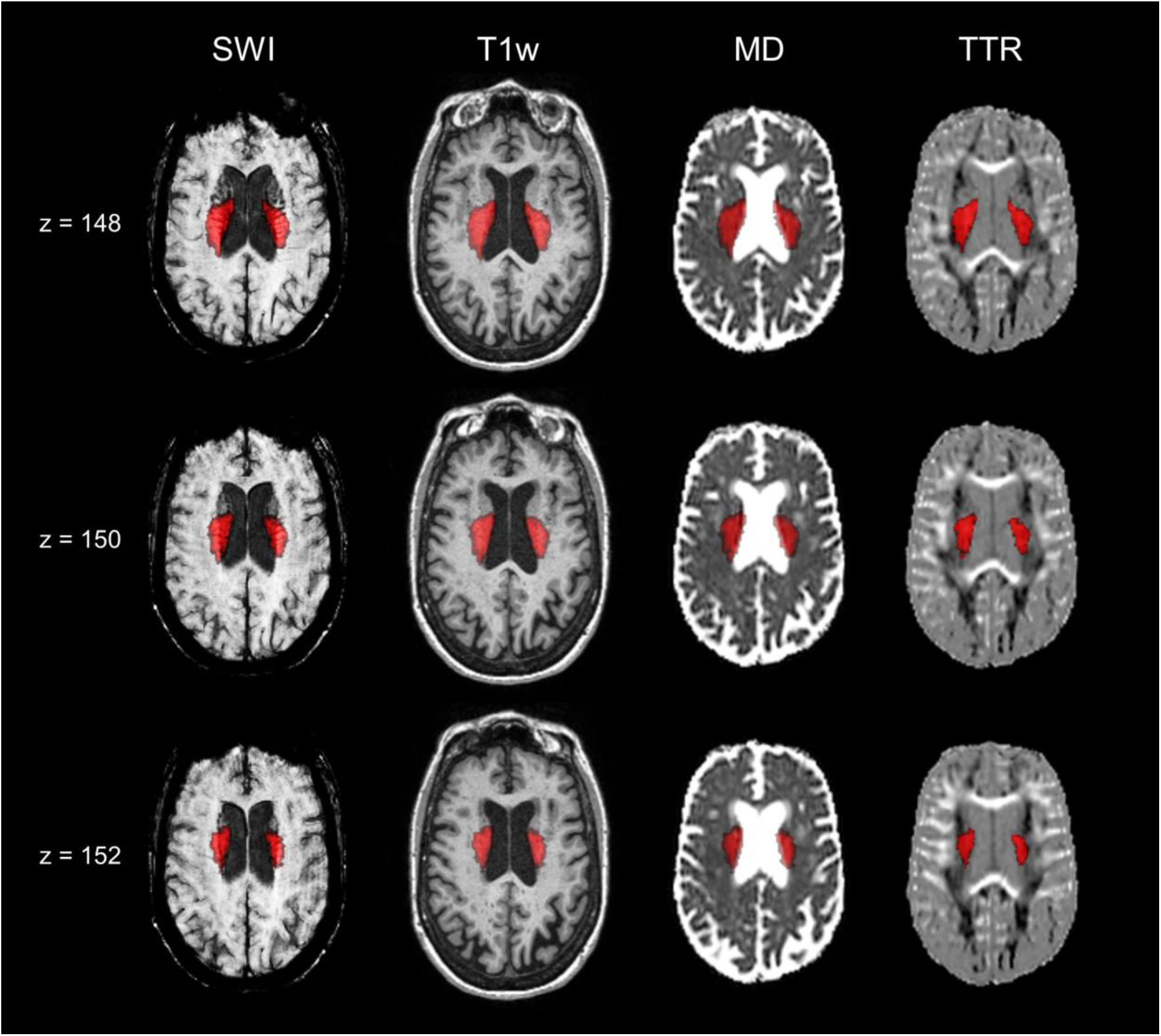
Automatically generated regions of interest (ROIs) for delineating periventricular regions based on mean diffusivity (MD) maps. The proposed region growth algorithm was optimized using multimodal imaging data, including susceptibility-weighted imaging (SWI), T1-weighted imaging (T1w), and MD maps. The resulting ROIs serve as sampling regions for transverse tensor ratio (TTR) maps, which is used to represent periventricular diffusivity.

### S4 Partial correlation analysis of DTI-ALPS and PVeD additionally adjusted for mean diffusivity

**Supplementary Figure 4.**
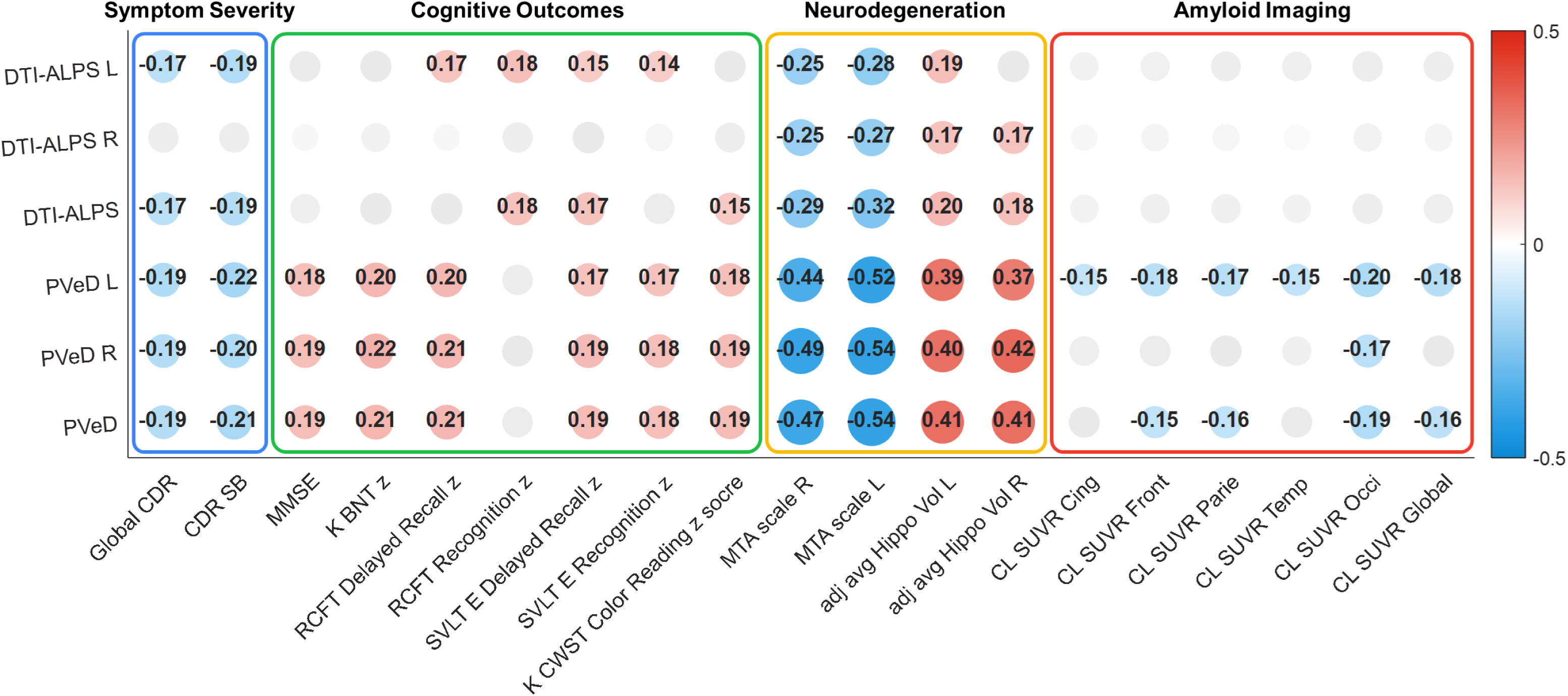
Partial correlation matrix of imaging-derived metrics with multifaceted clinical characteristics, with the additional statistical adjustment for mean diffusivity in the BICWALZS cohort. Imaging-derived metrics refer to the diffusion tensor imaging (DTI)-derived measures including DTI-ALPS and periventricular diffusivity (PVeD). The multifaceted clinical characteristics include four domains; symptom severity, cognitive outcomes, neurodegeneration, and Aβ burden. The matrix was generated using the BICWALZS dataset (n = 440). Values are Spearman’s correlation coefficients for each pairwise correlation. Covariates including age, sex, education, and mean diffusivity were adjusted. Hippocampal volume was additionally adjusted by the total intracranial volume. Darker blue and red colors indicate greater negative and positive correlations, respectively. Gray dots indicate those did not pass the significance threshold adjusted by multiple comparisons. Abbreviations: adj, adjusted; avg, average; CDR, Clinical Dementia Rating; CDR-SB, Clinical Dementia Rating Sum of Boxes; Cing, cingulum; CL, centiloid; CU, cognitively unimpaired; DTI, diffusion tensor image; DTI-ALPS, diffusion tensor image analysis along the perivascular space; Front, frontal; Hippo, hippocampal; K-BNT, Korean version of the Boston Naming Test; K-CWST, Korean version of the Color Word Stroop Test; L, left; MCI, mild cognitive impairment; MMSE, Mini-Mental State Examination; MTA, medial temporal lobe atrophy; Occi, occipital; Parie, parietal; PET, positron emission tomography; PVeD, periventricular diffusivity; R, right; RCFT, Rey Complex Figure Test and Recognition Trial; SUVR, standard uptake value ratio; SVLT-E, Seoul Verbal Learning Test Elderly’s version; Temp, temporal; Vol, volume; WM, white matter; z, z-score.

### S5 Interaction of DTI-ALPS with APOE4 on amyloid deposition

**Supplementary Figure 5.**
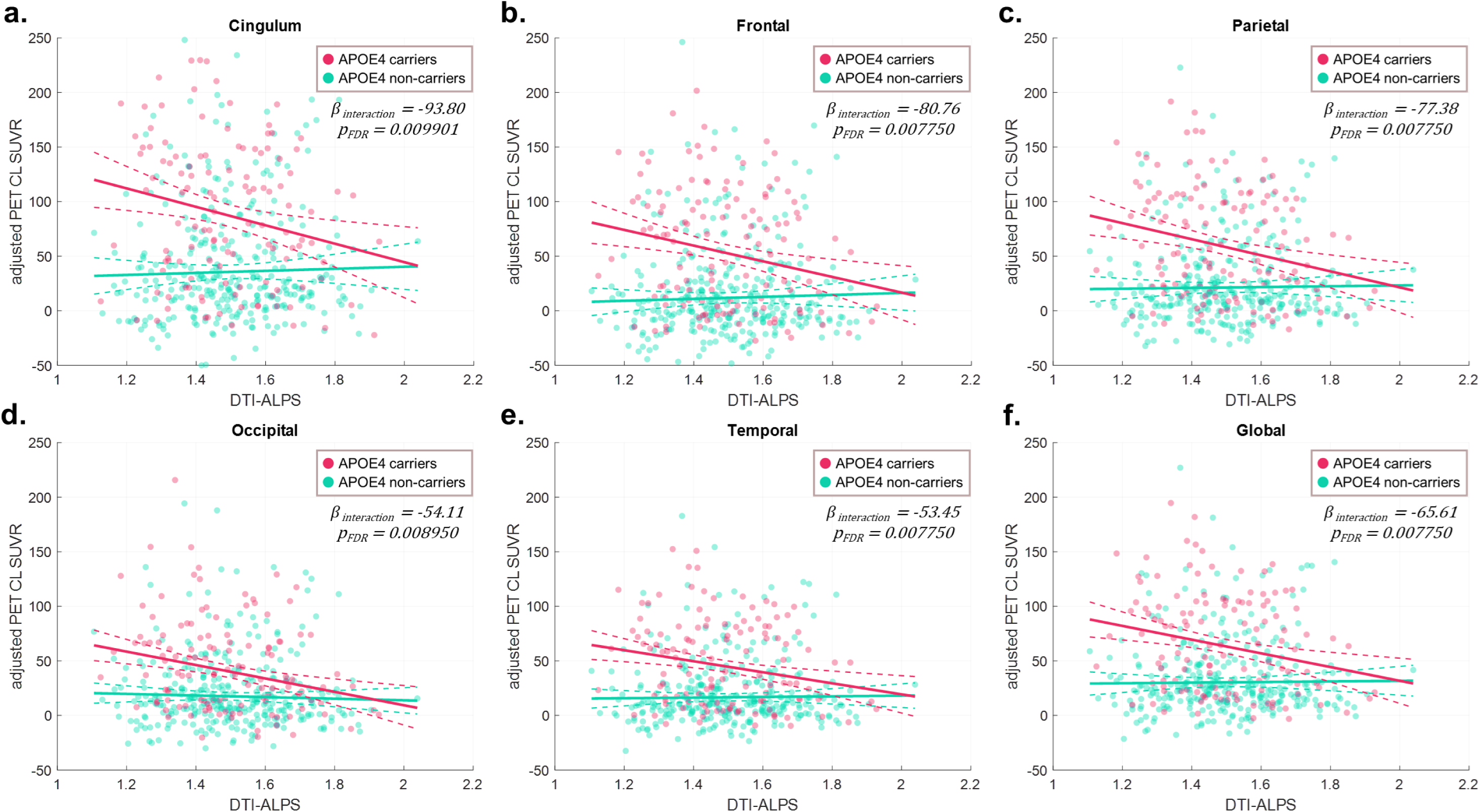
The presence of APOE4 allele enhances the association between DTI-ALPS and Aβ deposition in the BICWALZS cohort. Details of the statistical tests used are provided in Supplementary Table 5. The amyloid PET SUVRs were sampled from five anatomical regions including the cingulum (a), frontal lobe (b), parietal lobe (c), occipital lobe (d), and temporal lobe (e), with the inclusion of the global measure (f). The models included age, sex, and education as covariates. Multiple comparisons were controlled using the Benjamini-Hochberg correction accounting for the number of image measures analyzed. Green and red colors represent APOE4 non-carriers and APOE4 carriers, respectively. The interaction terms between DTI-ALPS and APOE4 are reported. Abbreviations: APOE, apolipoprotein E; CL, centiloid; DTI-ALPS, diffusion tensor image analysis along the perivascular space; FDR, false discovery rate; PET, positron emission tomography; SUVR, standard uptake value ratio.

### S6 Group comparisons of DTI-ALPS and PVeD between CU, MCI, and dementia

**Supplementary Figure 6.**
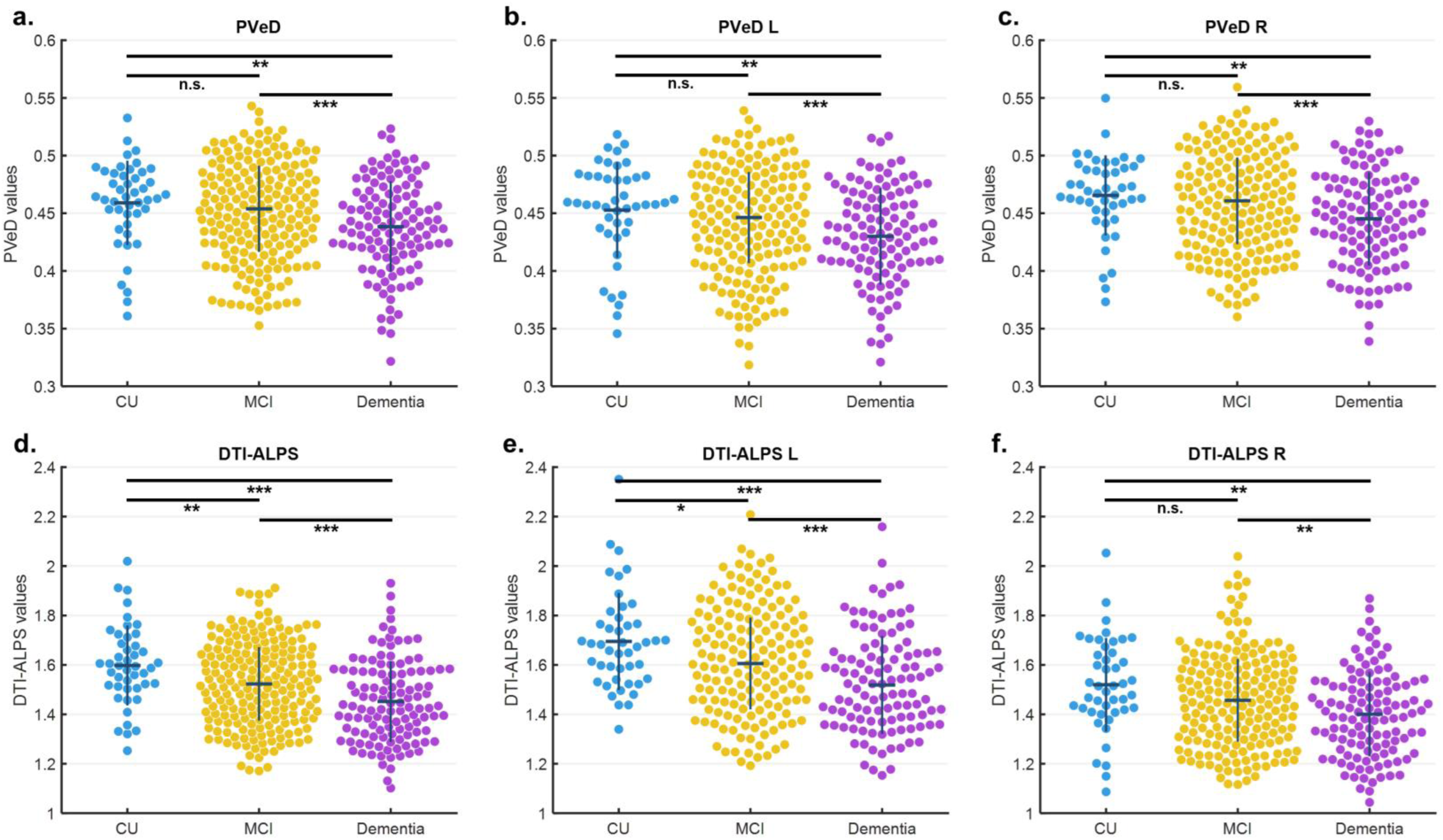
Beeswarm plots of PVeD and DTI-ALPS measures for group comparison in the BICWALZS cohort. Details of the statistical tests used are provided in Supplementary Table 7. Diffusion imaging-derived metrics for group comparisons include mean PVeD (a), left PVeD (b), right PVeD (c), mean DTI-ALPS index (d), left DTI-ALPS index (e), and right DTI-ALPS index (f).

### S7 Additional results in the replication cohort: the MCSA dataset

**Supplementary Figure 7.**
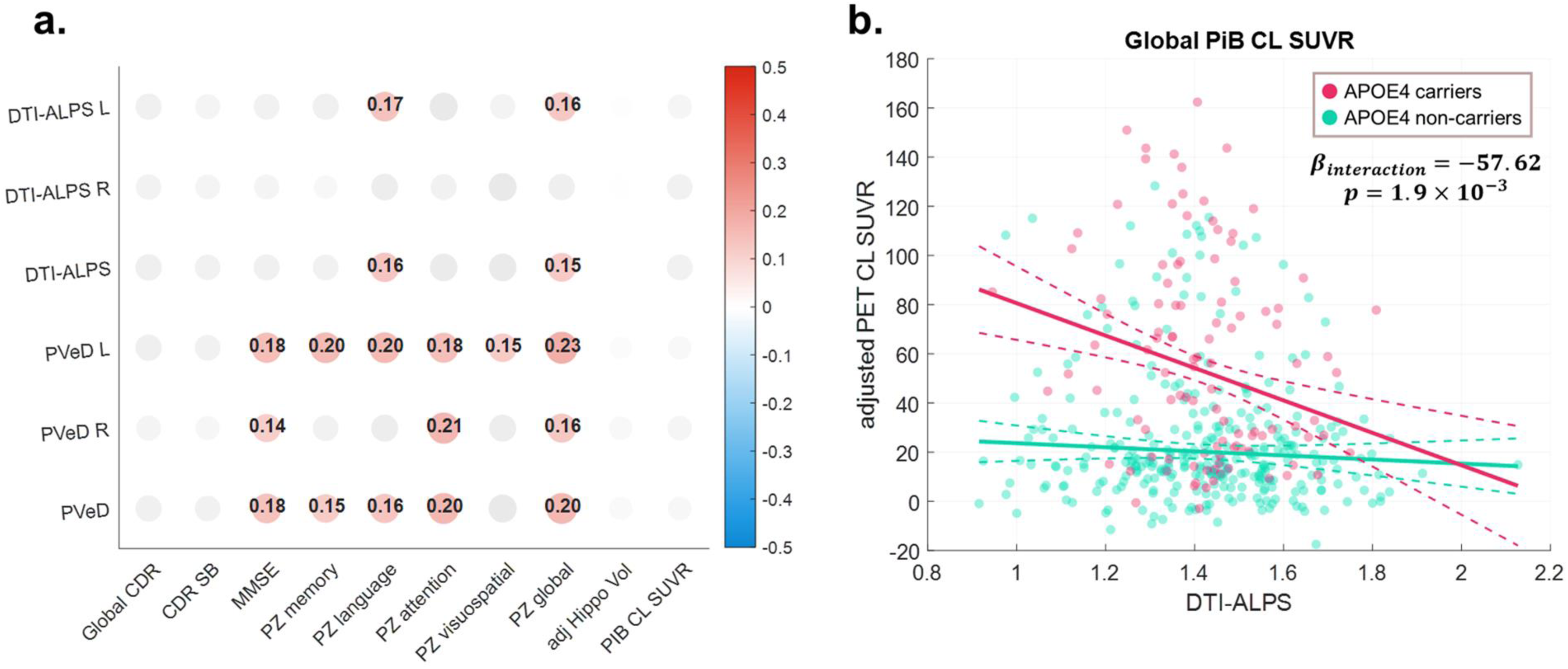
Partial correlation matrix of imaging-derived metrics with multifaceted clinical characteristics, with the additional statistical adjustment for mean diffusivity in the MCSA cohort (a), and the presence of APOE4 allele enhances the association between DTI-ALPS and Aβ deposition in the MCSA cohort (b).

**Supplementary Figure 8.**
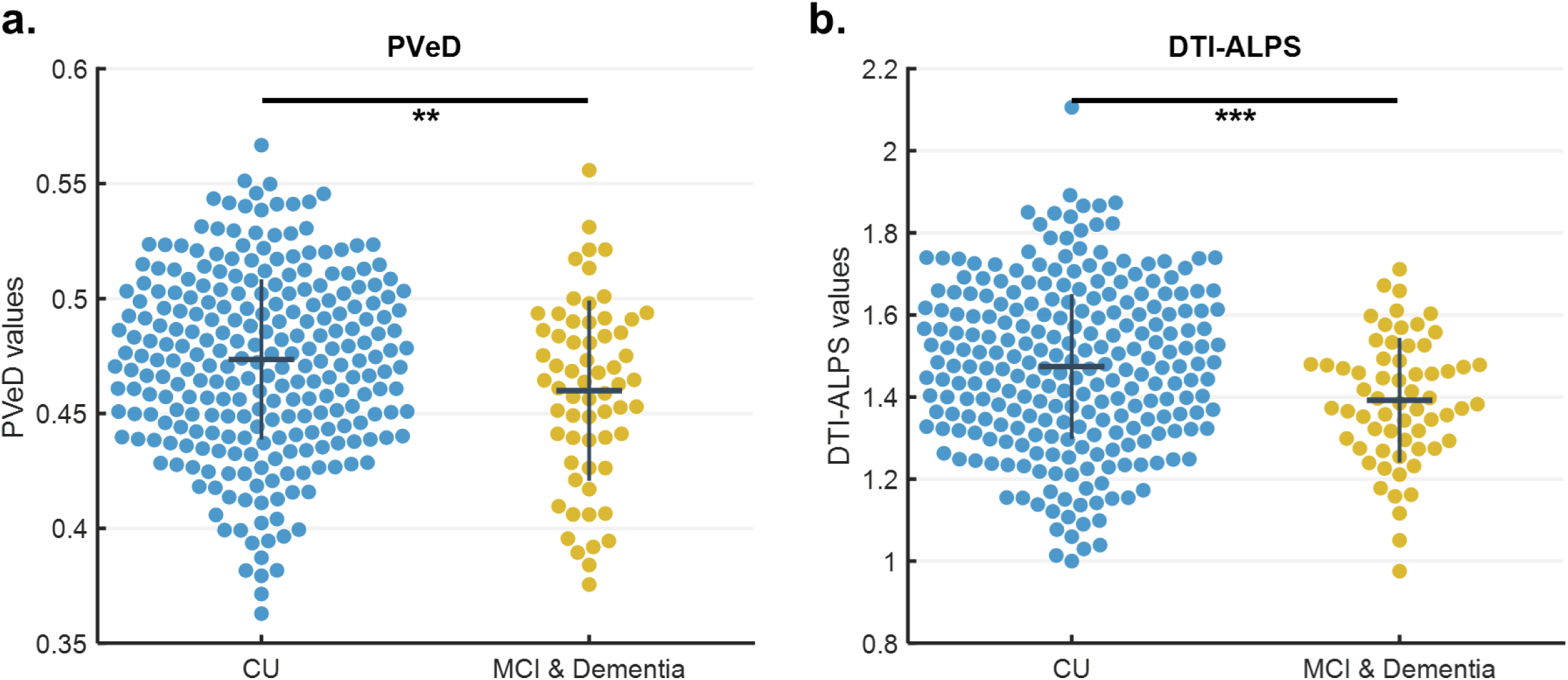
Beeswarm plots of PVeD (a) and DTI-ALPS (b) measures for group comparison in the MCSA cohort.

## Supplementary Tables

S. Table 1 Neuroimaging data acquisition parameters

S. Table 2 Demographic information of subjects in the MCSA replication cohort

S. Table 3 Partial correlation analysis of diffusion imaging-derived measures

S. Table 4 Interaction effects between PVeD and APOE4 on amyloid deposition

S. Table 5 Interaction effects between DTI-ALPS and APOE4 on amyloid deposition

S. Table 6 Association between baseline PVeD and longitudinal cognitive changes

S. Table 7 Group comparisons of DTI-ALPS and PVeD

S. Table 8 Additional results in the replication cohort: the MCSA dataset

**Supplementary Table 1.**
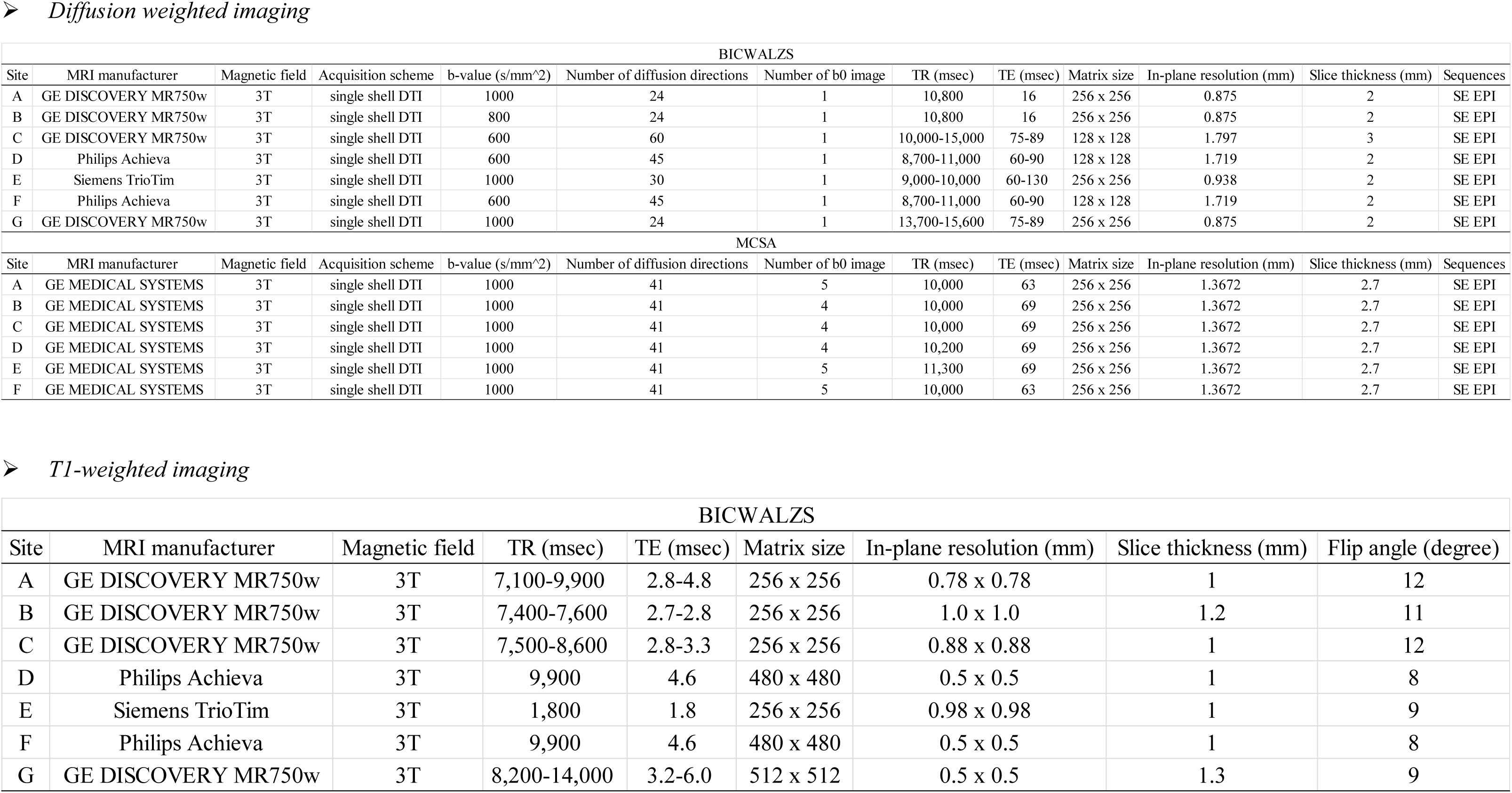
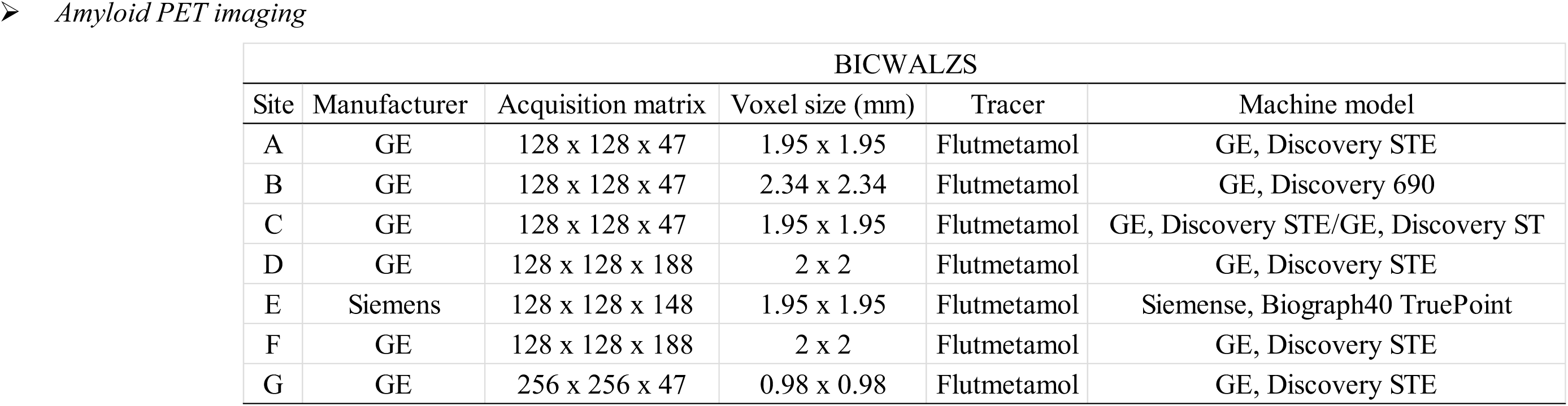
MRI, DTI, and PET acquisition parameters for the BICWALZS and DTI acquisition parameters for the MCSA.

➤ *Diffusion weighted imaging*
| BICWALZS |  |  |  |  |  |  |  |  |  |  |  |  |
| --- | --- | --- | --- | --- | --- | --- | --- | --- | --- | --- | --- | --- |
| Site | MRI manufacturer | Magnetic field | Acquisition scheme | b-value (s/mm^2) | Number of diffusion directions | Number of b0 image | TR (msec) | TE (msec) | Matrix size | In-plane resolution (mm) | Slice thickness (mm) | Sequences |
| A | GE DISCOVERY MR750w | 3T | single shell DTI | 1000 | 24 | 1 | 10,800 | 16 | 256 x 256 | 0.875 | 2 | SE EPI |
| B | GE DISCOVERY MR750w | 3T | single shell DTI | 800 | 24 | 1 | 10,800 | 16 | 256 x 256 | 0.875 | 2 | SE EPI |
| C | GE DISCOVERY MR750w | 3T | single shell DTI | 600 | 60 | 1 | 10,000-15,000 | 75-89 | 128 x 128 | 1.797 | 3 | SE EPI |
| D | Philips Achieva | 3T | single shell DTI | 600 | 45 | 1 | 8,700-11,000 | 60-90 | 128 x 128 | 1.719 | 2 | SE EPI |
| E | Siemens TrioTim | 3T | single shell DTI | 1000 | 30 | 1 | 9,000-10,000 | 60-130 | 256 x 256 | 0.938 | 2 | SE EPI |
| F | Philips Achieva | 3T | single shell DTI | 600 | 45 | 1 | 8,700-11,000 | 60-90 | 128 x 128 | 1.719 | 2 | SE EPI |
| G | GE DISCOVERY MR750w | 3T | single shell DTI | 1000 | 24 | 1 | 13,700-15,600 | 75-89 | 256 x 256 | 0.875 | 2 | SE EPI |
| MCSA |  |  |  |  |  |  |  |  |  |  |  |  |
| Site | MRI manufacturer | Magnetic field | Acquisition scheme | b-value (s/mm^2) | Number of diffusion directions | Number of b0 image | TR (msec) | TE (msec) | Matrix size | In-plane resolution (mm) | Slice thickness (mm) | Sequences |
| A | GE MEDICAL SYSTEMS | 3T | single shell DTI | 1000 | 41 | 5 | 10,000 | 63 | 256 x 256 | 1.3672 | 2.7 | SE EPI |
| B | GE MEDICAL SYSTEMS | 3T | single shell DTI | 1000 | 41 | 4 | 10,000 | 69 | 256 x 256 | 1.3672 | 2.7 | SE EPI |
| C | GE MEDICAL SYSTEMS | 3T | single shell DTI | 1000 | 41 | 4 | 10,000 | 69 | 256 x 256 | 1.3672 | 2.7 | SE EPI |
| D | GE MEDICAL SYSTEMS | 3T | single shell DTI | 1000 | 41 | 4 | 10,200 | 69 | 256 x 256 | 1.3672 | 2.7 | SE EPI |
| E | GE MEDICAL SYSTEMS | 3T | single shell DTI | 1000 | 41 | 5 | 11,300 | 69 | 256 x 256 | 1.3672 | 2.7 | SE EPI |
| F | GE MEDICAL SYSTEMS | 3T | single shell DTI | 1000 | 41 | 5 | 10,000 | 63 | 256 x 256 | 1.3672 | 2.7 | SE EPI |

➤ *T1-weighted imaging*
| BICWALZS |  |  |  |  |  |  |  |  |
| --- | --- | --- | --- | --- | --- | --- | --- | --- |
| Site | MRI manufacturer | Magnetic field | TR (msec) | TE (msec) | Matrix size | In-plane resolution (mm) | Slice thickness (mm) | Flip angle (degree) |
| A | GE DISCOVERY MR750w | 3T | 7,100-9,900 | 2.8-4.8 | 256 x 256 | 0.78 x 0.78 | 1 | 12 |
| B | GE DISCOVERY MR750w | 3T | 7,400-7,600 | 2.7-2.8 | 256 x 256 | 1.0 x 1.0 | 1.2 | 11 |
| C | GE DISCOVERY MR750w | 3T | 7,500-8,600 | 2.8-3.3 | 256 x 256 | 0.88 x 0.88 | 1 | 12 |
| D | Philips Achieva | 3T | 9,900 | 4.6 | 480 x 480 | 0.5 x 0.5 | 1 | 8 |
| E | Siemens TrioTim | 3T | 1,800 | 1.8 | 256 x 256 | 0.98 x 0.98 | 1 | 9 |
| F | Philips Achieva | 3T | 9,900 | 4.6 | 480 x 480 | 0.5 x 0.5 | 1 | 8 |
| G | GE DISCOVERY MR750w | 3T | 8,200-14,000 | 3.2-6.0 | 512 x 512 | 0.5 x 0.5 | 1.3 | 9 |

➤ *Amyloid PET imaging*
| BICWALZS |  |  |  |  |  |
| --- | --- | --- | --- | --- | --- |
| Site | Manufacturer | Acquisition matrix | Voxel size (mm) | Tracer | Machine model |
| A | GE | 128 x 128 x 47 | 1.95 x 1.95 | Flutmetamol | GE, Discovery STE |
| B | GE | 128 x 128 x 47 | 2.34 x 2.34 | Flutmetamol | GE, Discovery 690 |
| C | GE | 128 x 128 x 47 | 1.95 x 1.95 | Flutmetamol | GE, Discovery STE/GE, Discovery ST |
| D | GE | 128 x 128 x 188 | 2 x 2 | Flutmetamol | GE, Discovery STE |
| E | Siemens | 128 x 128 x 148 | 1.95 x 1.95 | Flutmetamol | Siemens, Biograph40 TruePoint |
| F | GE | 128 x 128 x 188 | 2 x 2 | Flutmetamol | GE, Discovery STE |
| G | GE | 256 x 256 x 47 | 0.98 x 0.98 | Flutmetamol | GE, Discovery STE |

**Supplementary Table 2.** Characteristics of the replication cohort from MCSA.

| Demographic characteristics | CU | MCI & Dementia | Comparisons |
| --- | --- | --- | --- |
| N | 351 | 63 | - |
| Age, year | 74.7 ($\pm 8.5$) | 79.1 ($\pm 7.0$) | $p < 0.001$ |
| Sex, female, N (%) | 168 (48.0%) | 21 (33.3%) | $p = 0.031$ |
| Education, year | 14.5 ($\pm 2.7$) | 13.9 ($\pm 3.0$) | $p = 0.125$ |
| MMSE | 28.3 ($\pm 1.2$) | 25.5 ($\pm 2.4$) | $p < 0.001$ |
| Global CDR | 0.0 ($\pm 0.1$) | 0.4 ($\pm 0.3$) | $p < 0.001$ |
| CDR SB | 0.0 ($\pm 0.2$) | 1.3 ($\pm 1.5$) | $p < 0.001$ |
| APOE4 +, N (%) | 83 (23.7%) | 23 (36.5%) | $p = 0.036$ |
| Neuropsychological characteristics |  |  |  |
| Population z-score memory | -0.028 ($\pm 1.000$) | -1.869 ($\pm 0.994$) | $p < 0.001$ |
| Population z-score language | -0.096 ($\pm 0.966$) | -1.399 ($\pm 1.265$) | $p < 0.001$ |
| Population z-score attention | -0.113 ($\pm 0.957$) | -1.368 ($\pm 1.263$) | $p < 0.001$ |
| Population z-score visuospatial | 0.077 ($\pm 0.992$) | -0.754 ($\pm 0.964$) | $p < 0.001$ |
| Population z-score global | -0.037 ($\pm 0.935$) | -1.646 ($\pm 0.873$) | $p < 0.001$ |
| Imaging characteristics |  |  |  |
| Adjusted bilateral hippocampal volume | 7.296 ($\pm 0.745$) | 6.772 ($\pm 1.005$) | $p = 0.001$ |
| Amyloid PET PiB global CL SUVR | 28.573 ($\pm 32.343$) | 60.220 ($\pm 47.745$) | $p < 0.001$ |
| DTI-ALPS | 1.474 ($\pm 0.176$) | 1.392 ($\pm 0.152$) | $p < 0.001$ |
| PVeD | 0.474 ($\pm 0.035$) | 0.460 ($\pm 0.039$) | $p = 0.007$ |

1. Comparisons were adjusted for age, sex, and education in the neuropsychological and imaging measures
2. Hippocampal volume was adjusted by total intracranial volume
Abbreviations: APOE, apolipoprotein E; CDR, Clinical Dementia Rating; CDR-SB, Clinical Dementia Rating
Sum of Boxes; CL, centiloid; CU, cognitively unimpaired; DTI-ALPS, diffusion tensor image analysis along
the perivascular space; MCI, mild cognitive impairment; MMSE, Mini-Mental State Examination; PET,
positron emission tomography; PiB, Pittsburgh Compound B; PVeD, periventricular diffusivity; SUVR,
standard uptake value ratio.

**Supplementary Table 3.** Partial correlation analysis of diffusion imaging-derived measures.

➤ *Partial correlation coefficients in Figure 3a.*
|  | Global CDR | CDR SB | MMSE | K BNT z | RCFT Delayed Recall z | RCFT Recognition z | SVLT E Delayed Recall z | SVLT E Recognition z | K CWST Color Reading z socre |
| --- | --- | --- | --- | --- | --- | --- | --- | --- | --- |
| DTI-ALPS | -0.21067<br>(8.94E-06) | -0.19071<br>(6.02E-05) | 0.15671<br>(1.01E-03) | 0.15645<br>(2.80E-03) | 0.15349<br>(1.37E-03) | 0.16117<br>(8.18E-04) | 0.16481<br>(5.42E-04) | 0.12786<br>(7.45E-03) | 0.22191<br>(4.88E-06) |
| PVeD | -0.17251<br>(2.91E-04) | -0.19571<br>(3.80E-05) | 0.17032<br>(3.48E-04) | 0.20663<br>(7.30E-05) | 0.19603<br>(4.09E-05) | 0.11531<br>(1.70E-02) | 0.18250<br>(1.25E-04) | 0.17476<br>(2.41E-04) | 0.16337<br>(8.24E-04) |
| WM FA | -0.21184<br>(7.95E-06) | -0.16898<br>(3.88E-04) | 0.19223<br>(5.24E-05) | 0.15223<br>(3.64E-03) | 0.15883<br>(9.24E-04) | 0.12969<br>(7.22E-03) | 0.12990<br>(6.54E-03) | 0.09327<br>(5.14E-02) | 0.24649<br>(3.56E-07) |
| WM MD | 0.27423<br>(5.60E-09) | 0.27165<br>(7.87E-09) | -0.26566<br>(1.71E-08) | -0.22717<br>(1.24E-05) | -0.24123<br>(3.89E-07) | -0.19666<br>(4.19E-05) | -0.21435<br>(6.16E-06) | -0.17454<br>(2.46E-04) | -0.29799<br>(5.61E-10) |

|  | MTA scale L | MTA scale R | adj avg Hippo Vol L | adj avg Hippo Vol R | CL SUVR Cing | CL SUVR Front | CL SUVR Parie | CL SUVR Temp | CL SUVR Occi | CL SUVR Global |
| --- | --- | --- | --- | --- | --- | --- | --- | --- | --- | --- |
| DTI-ALPS | -0.36897<br>(1.53E-15) | -0.33793<br>(3.91E-13) | 0.23112<br>(1.17E-06) | 0.18070<br>(1.57E-04) | -0.02783<br>(5.62E-01) | -0.05675<br>(2.36E-01) | -0.07589<br>(1.13E-01) | -0.05333<br>(2.66E-01) | -0.10901<br>(2.27E-02) | -0.07477<br>(1.19E-01) |
| PVeD | -0.51149<br>(1.67E-30) | -0.45025<br>(3.35E-23) | 0.39028<br>(3.33E-17) | 0.40096<br>(3.74E-18) | -0.13306<br>(5.34E-03) | -0.15557<br>(1.10E-03) | -0.15539<br>(1.12E-03) | -0.12641<br>(8.16E-03) | -0.18715<br>(8.29E-05) | -0.15970<br>(8.07E-04) |
| WM FA | -0.33181<br>(1.09E-12) | -0.34649<br>(9.01E-14) | 0.25865<br>(4.77E-08) | 0.21065<br>(9.85E-06) | 0.00843<br>(8.61E-01) | 0.00078<br>(9.87E-01) | -0.03579<br>(4.55E-01) | -0.02899<br>(5.46E-01) | -0.08487<br>(7.64E-02) | -0.03322<br>(4.89E-01) |
| WM MD | 0.52663<br>(1.50E-32) | 0.53456<br>(1.16E-33) | -0.46818<br>(5.67E-25) | -0.43703<br>(1.27E-21) | 0.04226<br>(3.78E-01) | 0.04962<br>(3.01E-01) | 0.06885<br>(1.51E-01) | 0.07665<br>(1.10E-01) | 0.13837<br>(3.75E-03) | 0.08132<br>(8.95E-02) |

\* Values in cells: Correlation (raw p-values)
\*\* Corrected Significance Threshold = 0.0031

➤ *Partial correlation coefficients in Figure 3b.*
|  | Global CDR | CDR SB | MMSE | K BNT z | RCFT Delayed Recall z | RCFT Recognition z | SVLT E Delayed Recall z | SVLT E Recognition z | K CWST Color Reading z score |
| --- | --- | --- | --- | --- | --- | --- | --- | --- | --- |
| DTI-ALPS L | -0.20967<br>(9.88E-06) | -0.19646<br>(3.54E-05) | 0.17247<br>(2.92E-04) | 0.15180<br>(3.74E-03) | 0.17919<br>(1.81E-04) | 0.16532<br>(5.95E-04) | 0.14562<br>(2.28E-03) | 0.14612<br>(2.20E-03) | 0.20433<br>(2.68E-05) |
| DTI-ALPS R | -0.15907<br>(8.47E-04) | -0.12775<br>(7.50E-03) | 0.10293<br>(3.14E-02) | 0.11378<br>(3.02E-02) | 0.07916<br>(1.00E-01) | 0.11410<br>(1.82E-02) | 0.14087<br>(3.17E-03) | 0.07575<br>(1.14E-01) | 0.17130<br>(4.49E-04) |
| PVeD L | -0.18195<br>(1.31E-04) | -0.20977<br>(9.78E-06) | 0.16901<br>(3.87E-04) | 0.20270<br>(1.01E-04) | 0.19496<br>(4.51E-05) | 0.11389<br>(1.84E-02) | 0.16990<br>(3.61E-04) | 0.16864<br>(3.99E-04) | 0.15653<br>(1.36E-03) |
| PVeD R | -0.17277<br>(2.85E-04) | -0.18384<br>(1.11E-04) | 0.16996<br>(3.59E-04) | 0.21461<br>(3.74E-05) | 0.19567<br>(4.22E-05) | 0.12324<br>(1.07E-02) | 0.18647<br>(8.81E-05) | 0.17850<br>(1.76E-04) | 0.16822<br>(5.70E-04) |

|  | MTA scale L | MTA scale R | adj avg Hippo Vol L | adj avg Hippo Vol R | CL SUVR Cing | CL SUVR Front | CL SUVR Parie | CL SUVR Temp | CL SUVR Occi | CL SUVR Global |
| --- | --- | --- | --- | --- | --- | --- | --- | --- | --- | --- |
| DTI-ALPS L | -0.33435<br>(7.13E-13) | -0.30187<br>(1.17E-10) | 0.21664<br>(5.39E-06) | 0.14988<br>(1.76E-03) | -0.03671<br>(4.44E-01) | -0.05887<br>(2.19E-01) | -0.08824<br>(6.53E-02) | -0.07282<br>(1.29E-01) | -0.11322<br>(1.79E-02) | -0.08323<br>(8.22E-02) |
| DTI-ALPS R | -0.32089<br>(6.36E-12) | -0.29853<br>(1.91E-10) | 0.20355<br>(1.97E-05) | 0.17460<br>(2.61E-04) | -0.01002<br>(8.35E-01) | -0.04093<br>(3.93E-01) | -0.04757<br>(3.21E-01) | -0.02126<br>(6.58E-01) | -0.08017<br>(9.42E-02) | -0.04884<br>(3.08E-01) |
| PVeD L | -0.49154<br>(5.80E-28) | -0.41837<br>(6.05E-20) | 0.37936<br>(2.87E-16) | 0.36343<br>(5.75E-15) | -0.14992<br>(1.67E-03) | -0.17673<br>(2.05E-04) | -0.17286<br>(2.83E-04) | -0.15228<br>(1.41E-03) | -0.20003<br>(2.53E-05) | -0.17946<br>(1.62E-04) |
| PVeD R | -0.51027<br>(2.41E-30) | -0.46665<br>(5.14E-25) | 0.38586<br>(8.04E-17) | 0.41645<br>(1.36E-19) | -0.10816<br>(2.37E-02) | -0.12709<br>(7.82E-03) | -0.13584<br>(4.44E-03) | -0.09853<br>(3.95E-02) | -0.16771<br>(4.30E-04) | -0.13457<br>(4.83E-03) |

\* Values in cells: Correlation (raw p-values)
\*\* Corrected Significance Threshold = 0.0031

**Supplementary Table 4.** Interaction effects between PVeD and APOE4 status on amyloid deposition.

➤ *Interaction effects in Figure 5.*
|  | PVeD | APOE4 | PVeD*APOE4 |  | PVeD | APOE4 | PVeD*APOE4 |
| --- | --- | --- | --- | --- | --- | --- | --- |
| Coefficients | -46.16 | 281.34 | -518.73 | Coefficients | -52.12 | 172.34 | -336.66 |
| SE | 82.05 | 66.23 | 149.36 | SE | 45.42 | 36.67 | 82.69 |
| $p_{\text{FDR}}$ | 0.6889 | < 0.0001 | 0.0006 | $p_{\text{FDR}}$ | 0.6889 | < 0.0001 | 0.0001 |
| Response: CL SUVR cingulum |  |  |  | Response: CL SUVR occipital lobe |  |  |  |
|  | PVeD | APOE4 | PVeD*APOE4 |  | PVeD | APOE4 | PVeD*APOE4 |
| Coefficients | -40.60 | 226.06 | -415.26 | Coefficients | -8.07 | 167.41 | -313.70 |
| SE | 61.17 | 49.38 | 111.35 | SE | 42.71 | 34.47 | 77.74 |
| $p_{\text{FDR}}$ | 0.6889 | < 0.0001 | 0.0003 | $p_{\text{FDR}}$ | 0.8502 | < 0.0001 | 0.0001 |
| Response: CL SUVR frontal lobe |  |  |  | Response: CL SUVR temporal lobe |  |  |  |
|  | PVeD | APOE4 | PVeD*APOE4 |  | PVeD | APOE4 | PVeD*APOE4 |
| Coefficients | -48.50 | 231.94 | -437.88 | Coefficients | -37.24 | 195.30 | -365.45 |
| SE | 56.47 | 45.59 | 102.80 | SE | 50.70 | 40.92 | 92.28 |
| $p_{\text{FDR}}$ | 0.6889 | < 0.0001 | 0.0001 | $p_{\text{FDR}}$ | 0.6889 | < 0.0001 | 0.0001 |
| Response: CL SUVR parietal lobe |  |  |  | Response: CL SUVR global |  |  |  |

\* Other covariates included age, sex, and education

**Supplementary Table 5.** Interaction effects between DTI-ALPS and APOE4 status on amyloid deposition.

|  | DTI-ALPS | APOE4 | DTI-ALPS*APOE4 |  | DTI-ALPS | APOE4 | DTI-ALPS*APOE4 |
| --- | --- | --- | --- | --- | --- | --- | --- |
| Coefficients | 9.36 | 191.80 | -93.80 | Coefficients | -7.32 | 103.64 | -54.11 |
| SE | 20.02 | 54.31 | 36.21 | SE | 11.13 | 30.19 | 20.13 |
| $p_{FDR}$ | 0.8286 | 0.0005 | 0.0099 | $p_{FDR}$ | 0.8286 | 0.0007 | 0.009 |
| Response: CL SUVR cingulum |  |  |  | Response: CL SUVR occipital lobe |  |  |  |
|  | DTI-ALPS | APOE4 | DTI-ALPS*APOE4 |  | DTI-ALPS | APOE4 | DTI-ALPS*APOE4 |
| Coefficients | 8.92 | 161.79 | -80.76 | Coefficients | 2.91 | 108.12 | -53.45 |
| SE | 15.20 | 41.25 | 27.50 | SE | 10.51 | 28.52 | 19.02 |
| $p_{FDR}$ | 0.8286 | 0.0003 | 0.0077 | $p_{FDR}$ | 0.8286 | 0.0003 | 0.0077 |
| Response: CL SUVR frontal lobe |  |  |  | Response: CL SUVR temporal lobe |  |  |  |
|  | DTI-ALPS | APOE4 | DTI-ALPS*APOE4 |  | DTI-ALPS | APOE4 | DTI-ALPS*APOE4 |
| Coefficients | 3.76 | 152.94 | -77.38 | Coefficients | 2.76 | 131.23 | -65.61 |
| SE | 14.13 | 38.34 | 25.56 | SE | 12.74 | 34.57 | 23.05 |
| $p_{FDR}$ | 0.8286 | 0.0003 | 0.0077 | $p_{FDR}$ | 0.8286 | 0.0003 | 0.0077 |
| Response: CL SUVR parietal lobe |  |  |  | Response: CL SUVR global |  |  |  |

\* Other covariates included age, sex, and education

**Supplementary Table 6.** Association between baseline PVeD and longitudinal cognitive changes.

➤ *Regression analysis in Figure 6.*
|  | PVeD L | PVeD R | PVeD | DTI-ALPS L | DTI-ALPS R | DTI-ALPS |
| --- | --- | --- | --- | --- | --- | --- |
| Coefficients | 14.734 | 16.425 | 16.698 | 0.716 | 1.756 | 1.784 |
| SE | 4.781 | 5.512 | 5.515 | 1.327 | 1.170 | 1.436 |
| $p_{\text{FDR}}$ | 0.00785 | 0.00785 | 0.00785 | 0.59123 | 0.20669 | 0.26180 |

\*Response: MMSE annual change rate
\*\* Other covariates included age, sex, and education

**Supplementary Table 7.** Group comparisons of DTI-ALPS and PVeD between CU, MCI, and dementia.

|  | PVeD | PVeD L | PVeD R |
| --- | --- | --- | --- |
| CU | 0.459 (0.037) | 0.453 (0.041) | 0.465 (0.035) |
| MCI | 0.454 (0.037) | 0.446 (0.040) | 0.461 (0.038) |
| Dementia | 0.438 (0.039) | 0.430 (0.042) | 0.445 (0.040) |
| Main effect | $F_{(2,434)} = 8.14, P_{\text{FDR}} = 0.000376$ | $F_{(2,434)} = 8.10, P_{\text{FDR}} = 0.000376$ | $F_{(2,434)} = 8.03, P_{\text{FDR}} = 0.000376$ |
| CU vs. Dementia | $P_{\text{Bonferroni}} = 0.0054$ | $P_{\text{Bonferroni}} = 0.0056$ | $P_{\text{Bonferroni}} = 0.0059$ |
| CU vs. MCI | $P_{\text{Bonferroni}} = 1.0000$ | $P_{\text{Bonferroni}} = 1.0000$ | $P_{\text{Bonferroni}} = 1.0000$ |
| MCI vs. Dementia | $P_{\text{Bonferroni}} = 0.0007$ | $P_{\text{Bonferroni}} = 0.0008$ | $P_{\text{Bonferroni}} = 0.0007$ |
|  | DTI-ALPS | DTI-ALPS L | DTI-ALPS R |
| CU | 1.598 (0.162) | 1.695 (0.197) | 1.520 (0.189) |
| MCI | 1.523 (0.149) | 1.606 (0.186) | 1.457 (0.168) |
| Dementia | 1.452 (0.163) | 1.519 (0.198) | 1.400 (0.171) |
| Main effect | $F_{(2,434)} = 16.12, P_{\text{FDR}} < 0.000001$ | $F_{(2,434)} = 15.62, P_{\text{FDR}} < 0.000001$ | $F_{(2,434)} = 8.65, P_{\text{FDR}} = 0.000376$ |
| CU vs. Dementia | $P_{\text{Bonferroni}} < 0.0001$ | $P_{\text{Bonferroni}} < 0.0001$ | $P_{\text{Bonferroni}} = 0.0011$ |
| CU vs. MCI | $P_{\text{Bonferroni}} = 0.0084$ | $P_{\text{Bonferroni}} = 0.0105$ | $P_{\text{Bonferroni}} = 0.0869$ |
| MCI vs. Dementia | $P_{\text{Bonferroni}} < 0.0001$ | $P_{\text{Bonferroni}} = 0.0001$ | $P_{\text{Bonferroni}} = 0.0064$ |

\* Other covariates included age, sex, and education
\*\* Following a significant ANCOVA, pairwise comparisons were conducted using the Bonferroni method to examine differences between adjusted group
means while controlling for multiple comparisons.

**Supplementary Table 8.** Additional results in the replication cohort: the MCSA dataset.

➤ *Partial correlation coefficients in Figure 7a.*
|  | Global CDR | CDR SB | MMSE | PZ memory | PZ language | PZ attention | PZ visuospatial | PZ global | adj Hippo Vol | PIB CL SUVR |
| --- | --- | --- | --- | --- | --- | --- | --- | --- | --- | --- |
| DTI-ALPS | -0.1423<br>(3.93E-03) | -0.1183<br>(1.67E-02) | 0.1327<br>(7.35E-03) | 0.1058<br>(3.43E-02) | 0.2001<br>(6.08E-05) | 0.1419<br>(4.98E-03) | 0.1500<br>(2.94E-03) | 0.1831<br>(3.20E-04) | 0.0053<br>(9.14E-01) | -0.1094<br>(2.66E-02) |
| PVeD | -0.0897<br>(6.99E-02) | -0.0758<br>(1.26E-01) | 0.1645<br>(8.66E-04) | 0.1388<br>(5.36E-03) | 0.1552<br>(1.95E-03) | 0.1929<br>(1.26E-04) | 0.1326<br>(8.67E-03) | 0.1916<br>(1.65E-04) | -0.0425<br>(3.90E-01) | -0.0463<br>(3.49E-01) |
|  | Global CDR | CDR SB | MMSE | PZ memory | PZ language | PZ attention | PZ visuospatial | PZ global | adj Hippo Vol | PIB CL SUVR |
| DTI-ALPS L | -0.1411<br>(4.26E-03) | -0.1123<br>(2.32E-02) | 0.1303<br>(8.49E-03) | 0.1163<br>(1.98E-02) | 0.2108<br>(2.34E-05) | 0.1531<br>(2.43E-03) | 0.1081<br>(3.26E-02) | 0.1928<br>(1.50E-04) | 0.0176<br>(7.22E-01) | -0.0866<br>(7.97E-02) |
| DTI-ALPS R | -0.1076<br>(2.96E-02) | -0.0943<br>(5.68E-02) | 0.1007<br>(4.24E-02) | 0.0611<br>(2.22E-01) | 0.1334<br>(7.87E-03) | 0.1033<br>(4.15E-02) | 0.1495<br>(3.05E-03) | 0.1265<br>(1.33E-02) | 0.0064<br>(8.98E-01) | -0.1009<br>(4.10E-02) |
| PVeD L | -0.0952<br>(5.45E-02) | -0.0831<br>(9.33E-02) | 0.1778<br>(3.14E-04) | 0.1906<br>(1.23E-04) | 0.1913<br>(1.28E-04) | 0.1740<br>(5.55E-04) | 0.1445<br>(4.19E-03) | 0.2206<br>(1.35E-05) | -0.0265<br>(5.92E-01) | -0.0423<br>(3.93E-01) |
| PVeD R | -0.0652<br>(1.88E-01) | -0.0498<br>(3.15E-01) | 0.1271<br>(1.03E-02) | 0.0816<br>(1.03E-01) | 0.1056<br>(3.58E-02) | 0.1953<br>(1.04E-04) | 0.1114<br>(2.77E-02) | 0.1497<br>(3.37E-03) | -0.0490<br>(3.22E-01) | -0.0579<br>(2.42E-01) |

\* Values in cells: Correlation (raw p-values)
\*\* Corrected Significance Threshold = 0.0063

➤ *Interaction effects in Figure 7b.*
|  | PVeD | APOE4 | PVeD*APOE4 |
| --- | --- | --- | --- |
| Coefficients | 19.64 | 185.69 | -326.88 |
| SE | 38.92 | 44.29 | 95.09 |
| *p*-value | 0.6141 | < 0.0001 | 0.0006 |
| Response: PiB CL SUVR global |  |  |  |

\* Other covariates included age, sex, and education

➤ *Regression analysis in Figure 7d & 7e.*
|  | PVeD | DTI-ALPS |
| --- | --- | --- |
| Coefficients | 4.832 | 0.294 |
| SE | 1.618 | 0.334 |
| *p*<sub>FDR</sub> | 0.0066 | 0.3800 |

\*Response: MMSE annual change rate
\*\* Other covariates included age, sex, and education

